# The Arp1/11 Minifilament of Dynactin Primes the Endosomal Arp2/3 Complex

**DOI:** 10.1101/2020.05.29.123372

**Authors:** Artem I. Fokin, Violaine David, Ksenia Oguievetskaia, Emmanuel Derivery, Caroline E. Stone, Luyan Cao, Nathalie Rocques, Nicolas Molinie, Véronique Henriot, Magali Aumont-Nicaise, Maria-Victoria Hinckelmann, Frédéric Saudou, Christophe Le Clainche, Andrew P. Carter, Guillaume Romet-Lemonne, Alexis M. Gautreau

**Affiliations:** Laboratoire de Biologie Structurale de la Cellule, CNRS, Ecole Polytechnique, IP Paris, Palaiseau, France; Université Paris-Saclay, CEA, CNRS, Institute for Integrative Biology of the Cell (I2BC), Gif-sur-Yvette, France; MRC Laboratory of Molecular Biology, Cambridge, UK; Université de Paris, CNRS, Institut Jacques Monod, Paris, France; Univ. Grenoble Alpes, Inserm, U1216, CHU Grenoble Alpes, Grenoble Institut des Neurosciences, Grenoble, France

## Abstract

Dendritic actin networks develop from a first actin filament through branching by the Arp2/3 complex. At the surface of endosomes, the WASH complex activates the Arp2/3 complex and interacts with the Capping Protein for unclear reasons. Here we show that that the WASH complex interacts with Dynactin and uncaps it through its FAM21 subunit. In vitro, the uncapped Arp1/11 minifilament elongates an actin filament, which then primes the WASH-induced Arp2/3 branching reaction. In Dynactin-depleted cells or in cells where the WASH complex is reconstituted with a FAM21 mutant that cannot uncap Dynactin, formation of branched actin at the endosomal surface is impaired. Our results reveal the importance of the WASH complex in coordinating two complexes containing actin-related proteins.

**One Sentence Summary:** Dendritic actin networks grow in an autocatalytic manner starting from the uncapped minifilament of Dynactin.

## Main Text

Dendritic actin networks nucleated by the Actin-related protein (Arp) 2 and Arp3 containing complex exert pushing forces onto membranes (*1*). The WASH complex activates the Arp2/3 complex at the surface of endosomes and thereby promotes scission of transport intermediates (*2*, *3*). The WASH complex is associated with Capping Protein (CP) (*2*, *4*). CP is a freely diffusing heterodimer that interacts with the barbed end of actin filaments with high affinity and blocks their elongation (*5*). The FAM21 subunit of the WASH complex displays a conserved CP Interacting motif (CPI) in its extended tail (*6*). CPI motifs directly bind to CP and reduce its affinity for actin filaments through a conformational change (*6*, *7*). The interaction with CPI containing proteins is thought to bring CP to subcellular locations, where its activity is needed (*8*). Along this model, the WASH complex might thus recruit CP to restrict the growth of endosomal dendritic actin networks, which are indeed small, typically below the diffraction limit of optical microscopy (*9*).

Dynactin, the major activator of the microtubule motor Dynein, is a multiprotein complex that contains CP (*10*). Surprisingly for a microtubule-oriented machinery, Dynactin is organized around a minifilament containing one molecule of Arp11, one molecule of β-actin and eight molecules of Arp1 from the pointed to the barbed end (*11*). The barbed end composed of Arp1 is capped by CP. The WASH complex and Dynactin might be in proximity, since the WASH complex is recruited at the endosomal surface through the interaction of FAM21 with the VPS complex of the retromer (*12*–*14*) and that Dynactin interacts with sorting nexins of the retromer (*15*).

### Uncapping Dynactin

To examine the possibility that Dynactin interacts with the WASH complex, we used a transgenic mouse that expresses the Dynactin subunit DCTN2 fused to GFP in the brain (*16*). GFP immunoprecipitates from brain extract contained the WASH complex, indicating that the two complexes interact (Fig. 1A). To inactivate the CPI motif of FAM21, we introduced 3 point mutations in the residues conserved in all CPI containing proteins (*6*). The mutated CPI, CPI*, consists of L1024A, R1031A and P1040A. Compared to WT CPI, CPI* had a 400-fold decreased affinity for CP (Fig. 1B, fig. S1). The 3 mutations were introduced into full-length FAM21 and stable MCF10A cell lines expressing WT or mutant FAM21 were isolated. FAM21 CPI* did not affect the integration of FAM21 into the WASH complex, nor the proper localization of the WASH complex at the surface of endosomes, but reduced the association of WASH with CP to undetectable levels (Fig. 1C, fig. S2).

**Fig. 1.**
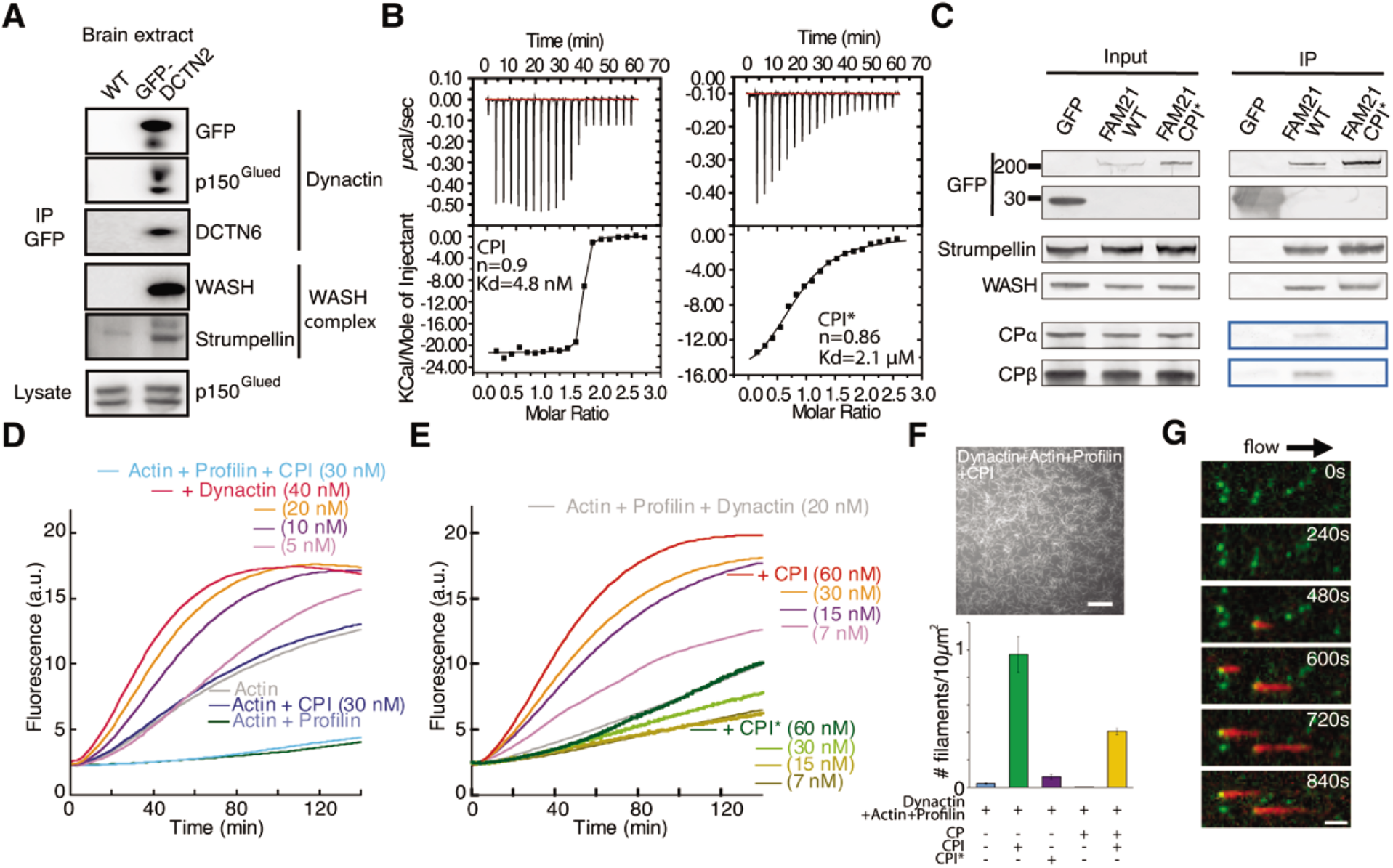
The CPI motif of FAM21 induces actin elongation from Dynactin. **A** The WASH complex coprecipitates with GFP tagged Dynactin (GFP-DCTN2) from the brain of a transgenic mouse. **B** Isothermal calorimetry of the interaction between CP and the wild type CPI or CPI*, a variant containing 3 point mutations. Curve fits indicate stoichiometry (n) and Kd of interaction. **C** Stable expression of GFP-FAM21 CPI* in MCF10A cells reconstitutes a WASH complex that does not recruit CP. **D-E** Pyrene-actin polymerization assays. Conditions: 2.0 μM actin (5 % labeled), 8 μM Profilin, native Dynactin, CPI or CPI*. n=3. **F** TIRF imaging of filaments elongated from surface anchored Dynactin in the presence of CPI. Observation 10 min after having introduced 1 μM labeled actin, 8.2 μM profilin, 0 or 2 μM CP, 0 or 2.7 μM CPI in F-buffer. Average and s.e.m. of 3 to 5 independent experiments performed with 2 independent preparations of GFP-Arp1 containing Dynactin. Scale bar: 20 μm. **G** Time-lapse images of actin filaments growing under flow from GFP-DCTN3 labeled Dynactin (green). Conditions: 1 μM actin (15% labeled, red), 1 μM Profilin, 50 nM CPI. Scale bar: 3 μm.

FAM21 CPI and CPI* had no activity on the polymerization of actin, but, as anticipated, CPI, but not CPI*, was able to release the inhibition of filament elongation due to CP, i.e. to ‘uncap’ actin filaments (fig. S3). We reasoned that if CPI was able to uncap Dynactin, actin elongation could ensue from the barbed end of the Arp1/11 minifilament. Indeed, Dynactin induced actin polymerization in the presence of CPI, but not in the presence of CPI* (Fig. 1, D and E). Profilin was added in this assay to suppress spontaneous actin nucleation (*17*). We next sought to visualize filament growth using TIRF microscopy. To this end, we purified GFP labeled Dynactin through tandem affinity purification and anchored it via the GFP binding protein to a passivated coverslip. When Dynactin was anchored, infusion of profilin-actin initiated the elongation of a few actin filaments (Fig. 1F). This basal activity of Dynactin was suppressed by the addition of purified CP, suggesting that some Dynactin complexes immunoprecipitated from cells were already uncapped. The number of elongating actin filaments was, however, significatively enhanced by CPI, but not by CPI*. Similar results were obtained with Dynactin was purified and anchored via GFP-Arp1 or via GFP-DCTN3 (Fig.1F, fig. S4A). The growth rate of elongating filaments unambiguously indicated that growing occurs through the barbed end (fig. S4B). We then used microfluidics and a sparse density of anchored GFP labeled Dynactin to monitor filament nucleation. Upon addition of CPI, the vast majority filaments elongated from a green dot corresponding to a single anchored Dynactin complex (Fig. 1G, fig. S4C,D). The rare occurrences where filaments did not elongate from a green dot (Fig. 4D) are likely to correspond to elongation from a bleached Dynactin complex, since filament fluctuations in the flow systematically indicated that filaments were specifically attached through an end (Movie S1). The simplest hypothesis that can account for all these observations is that Dynactin becomes uncapped by CPI.

To directly examine Dynactin uncapping, we took advantage of our dually tagged versions of Dynactin (His-mCherry-CPα with either Flag-GFP-Arp1 or Flag-GFP-DCTN3). Anchored Dynactin displayed high levels of colocalization of green and red spots, which were reduced upon incubation with CPI, but not CPI* (Fig. 2A). The single mCherry fluorophore was more prone to bleaching than the GFP (up to 8 copies for Arp1, up to 2 copies for DCTN3). This suggests that CPI dissociates CP from the rest of Dynactin. To unambiguously establish this point, we analyzed native Dynactin by gel filtration. Upon incubation with CPI, but not with CPI*, part of CP migrated in low molecular weight fractions and were thus no longer associated with the Dynactin peak (Fig. 2B; fig. S5 for full gels). Dynactin fractions incubated with CPI or not were negatively stained and imaged by electron microscopy. From the CPI treated sample, most single particles fell into two 3D classes that appeared to have lost density at the end of the rod-shaped structure (Fig. 2C, fig. S6). Docking a structural model of native Dynactin into the density map shows that Dynactin treated with CPI no longer accommodates CP at the barbed end of the Arp1/11 minifilament (Fig. 2D). Together these experiments demonstrate that Dynactin is uncapped when treated with CPI and that the uncapped Dynactin is a stable complex.

**Fig. 2.**
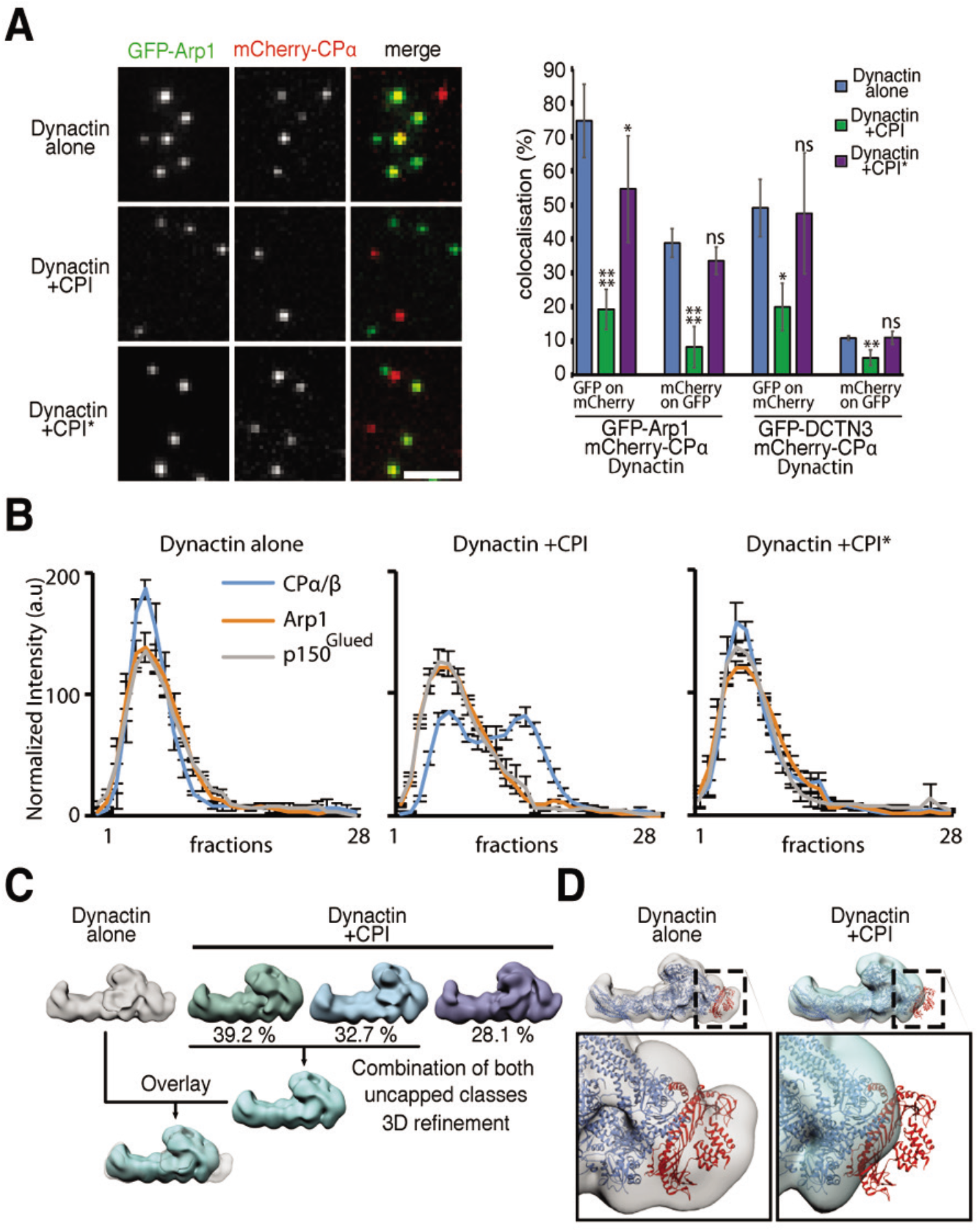
FAM21 CPI removes CP from the Arp1/11 minifilament of Dynactin. **A** Single Dynactin complexes containing GFP-Arp1 and mCherry-CPα were observed by TIRF microscopy. Quantification of the colocalization of green and red spots. Conditions: 40 nM of Dynactin preincubated with 2.7 μM CPI or CPI* for 1 min were diluted 10-fold and then adsorbed on the coverslip surface. Scale bar: 5 μm. Quantification of > 500 dots, 3 to 4 independent experiments with two different preparations of Dynactin purified through GFP-Arp1 and two others purified through GFP-DCTN3. Mean ± s.e.m., two-tailed Kruskal-Wallis test, * P<0.05, ** P<0.001, **** P<0.0001. **B** Distribution of Dynactin subunits in gel filtration in presence of CPI or CPI*. Mean ± s.e.m., n=3. **C** Elution fractions containing Dynactin were negatively stained and observed by electron microscopy (EM). Two major classes of 3D reconstructions obtained in the presence of CPI appear to lack a specific density. **D** The previously obtained cryo-EM model of Dynactin was fit into the negative stain EM densities. The two CP subunits are in red, other Dynactin subunits in blue.

### Arp2/3 Priming

The actin filament elongated from Dynactin might be useful to initiate an actin branching reaction. Indeed, the activated Arp2/3 complex nucleates a filament when it lands on a pre-existing filament (*18*). Since the actin filament is both a substrate and a product of the reaction, the generation of dendritic actin networks is an autocatalytic process that requires a first primer filament to get started (*19*). We verified that uncapped Dynactin can provide such a primer filament in a single filament TIRF assay. To this end, Dynactin was first uncapped by CPI in the presence of red actin and profilin. The resulting filaments were then mixed with the Arp2/3 complex, green actin and the Arp2/3 activating VCA motif of WASH. Numerous green filaments branched out from red actin filaments elongated from Dynactin (Fig. 3A). The density of actin branches was similar on filaments elongated from Dynactin (0.31 ± 0.11 branches/μm, n=15) and from spontaneously nucleated actin filaments (0.28 ± 0.12 branches/μm, n=19). Uncapped Dynactin is thus competent to initiate the formation of a branched actin structure.

**Fig. 3.**
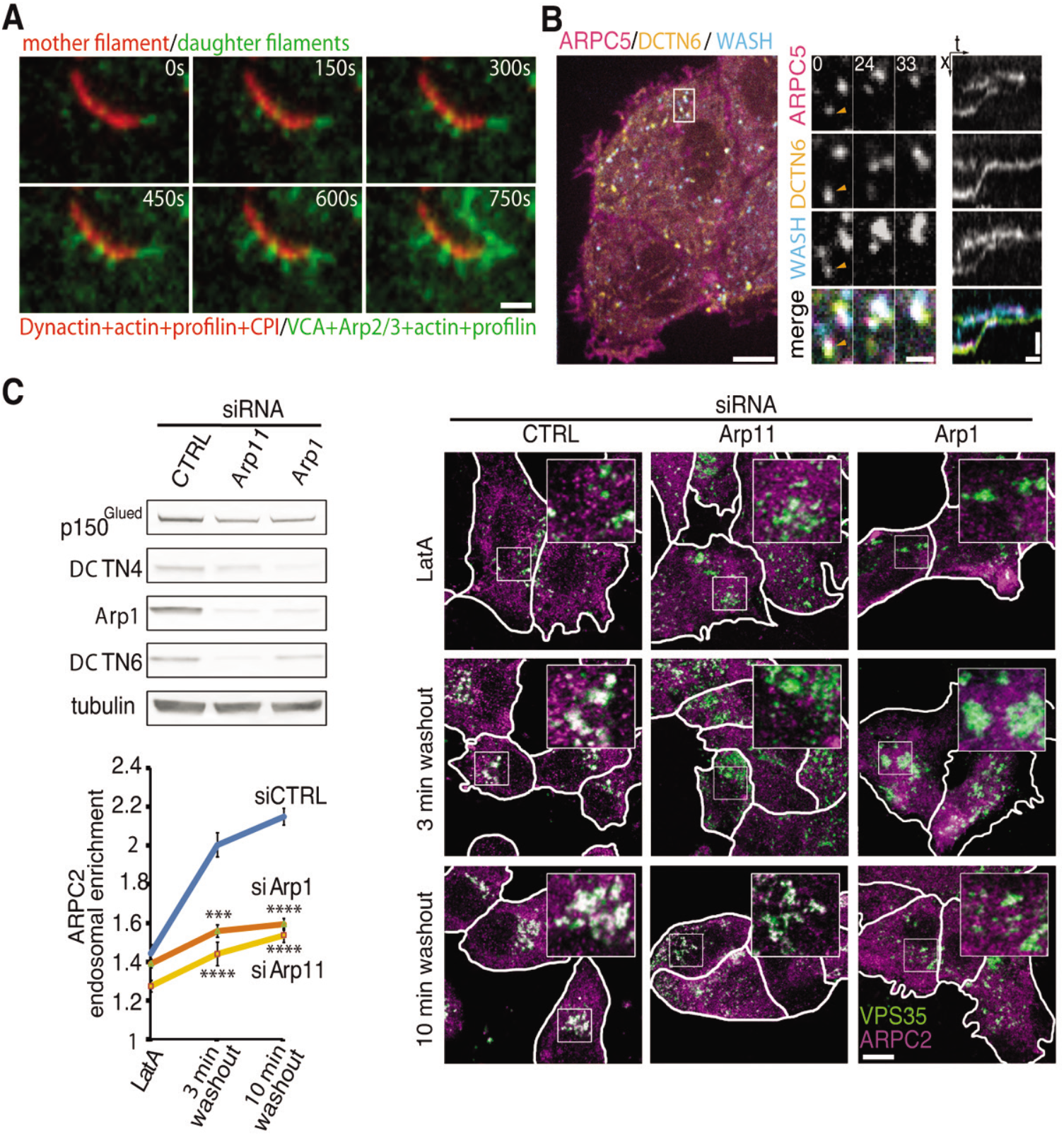
Dynactin primes endosomal dendritic actin networks. **A** Filaments elongated from Dynactin (2 nM Dynactin, 270 nM CPI, 1 μM profilin and 0.8 μM red Alexa568-actin) were diluted into (60 nM WASH VCA 60 nM Arp2/3, 0.4 μM profilin, 0.4 μM green Alexa 488-actin in F-buffer). The branching reaction is observed by TIRF microscopy. Scale bar: 2 μm. **B** Live spinning disk confocal microscopy of a MCF10A cell line stably expressing GFP-ARPC5 (Arp2/3), mCherry-DCTN6 (Dynactin) and iRFP-WASH. Single confocal plane. Scale bar: 6 μm. Middle panels: zoomed still images from the white box, elapsed time in seconds, scale bar: 1.2 μm. Right panels: kymograph, scale bars: vertical 1.2 μm, horizontal 12 s. **C** MCF10A cells were depleted of Dynactin through siRNAs targeting Arp1 or Arp11. Endosomal dendritic actin networks were estimated by the overlap of ARPC2 (Arp2/3) with VPS35 (retromer) upon LatrunculinA (LatA) washout. Scale bar: 10 μm. Mean ± s.e.m., two-tailed Kruskal–Wallis, *** P<0.001, **** P<0.0001. n=2.

The WASH complex activates the Arp2/3 complex at the surface of endosomes (*2*, *3*). To examine whether Dynactin was present in endosomal dendritic actin structures, we derived a triple transgenic MCF10A cell line stably expressing GFP-tagged ARPC5 (Arp2/3), mCherry-tagged DCTN6 (Dynactin) and iRFP-tagged WASH. In this cell line, fusion proteins are not present in excess and incorporate into their respective complexes as shown by sucrose gradient and immunoprecipitation (fig. S7A-C). We chose to tag DCTN6, because it was reported to be a subunit that targets Dynactin to endosomes (*20*). Using fast live cell imaging, DCTN6-tagged Dynactin exhibited the expected behavior of microtubule + end tracking (Movie S2). Importantly, some endosomal compartments transiently appeared simultaneously positive for WASH, Arp2/3 and Dynactin (Fig.3b; fig.7S, D and E). WASH and Arp2/3 spots were more often colocalized than WASH and Dynactin spots (59 ± 2 % *vs.* 6.2 ± 0.9 %, mean ± s.e.m., 1587 WASH spots in 28 cells). Kymograph analyses indicated that this association cannot be decomposed into a stereotypical sequence of events. Unlike clathrin coated pits, where branched actin polymerization is precisely coordinated with membrane scission (*21*), dendritic actin networks at the surface of endosomes perform several functions in addition to membrane scission (*9*, *22*, *23*).

Our in *vitro* reconstitution indicated that Dynactin should be required to initiate branched actin in the physiological presence of profilin. So we depleted Dynactin by targeting the minifilament forming subunits Arp1 and Arp11 and analyzed dendritic actin structures associated with retromer after wash-out of the actin depolymerizing drug latrunculin A. In this regrowth assay, Dynactin depletion significantly delayed the appearance of the Arp2/3 complex at the surface of endosomes (Fig. 3C). Even though endosomal WASH complexes could still in principle bind to the Arp2/3 complex, this result illustrates how formation of dendritic actin networks containing Arp2/3 at each branch is impaired upon Dynactin depletion. To our knowledge, this is the first example of actin polymerization that requires Dynactin. Since the depletion of Dynactin affects numerous cellular functions, we complemented this result by the phenotypic analysis of cells expressing FAM21 CPI*.

### Uncapping by WASH

When we knocked down endogenous FAM21 using siRNAs targeting the 3’UTR of FAM21, expression of FAM21 wild type or CPI* restored normal levels of FAM21 and of WASH to the surface of endosomes (fig. S8). In sharp contrast, however, only FAM21 wild type, and not FAM21 CPI*, was able to restore proper amounts of dendritic actin networks at the surface of endosomes (Fig. 4A). This experiment indicates that the interaction of the WASH complex with CP does not restrict the growth of endosomal branched actin. Rather, the ability of the WASH complex to uncap Dynactin is critical to generate dendritic actin networks at the surface of endosomes. We next examined the cell functions ascribed to the WASH complex in this system, where WASH complexes are present, but deficient in polymerizing branched actin. Using live imaging of cells loaded with fluorescent transferrin, we observed that knock-down of FAM21 led an increase in endosome size and tubulation (Fig. 4B), as previously described (*2*, *3*, *9*). These defects indicate an impaired scission of the transport intermediates that sort endosomal cargoes (*23*). We found that FAM21 wild type, but not CPI*, was able to rescue the morphological defects induced by FAM21 knock-down.

**Fig. 4.**
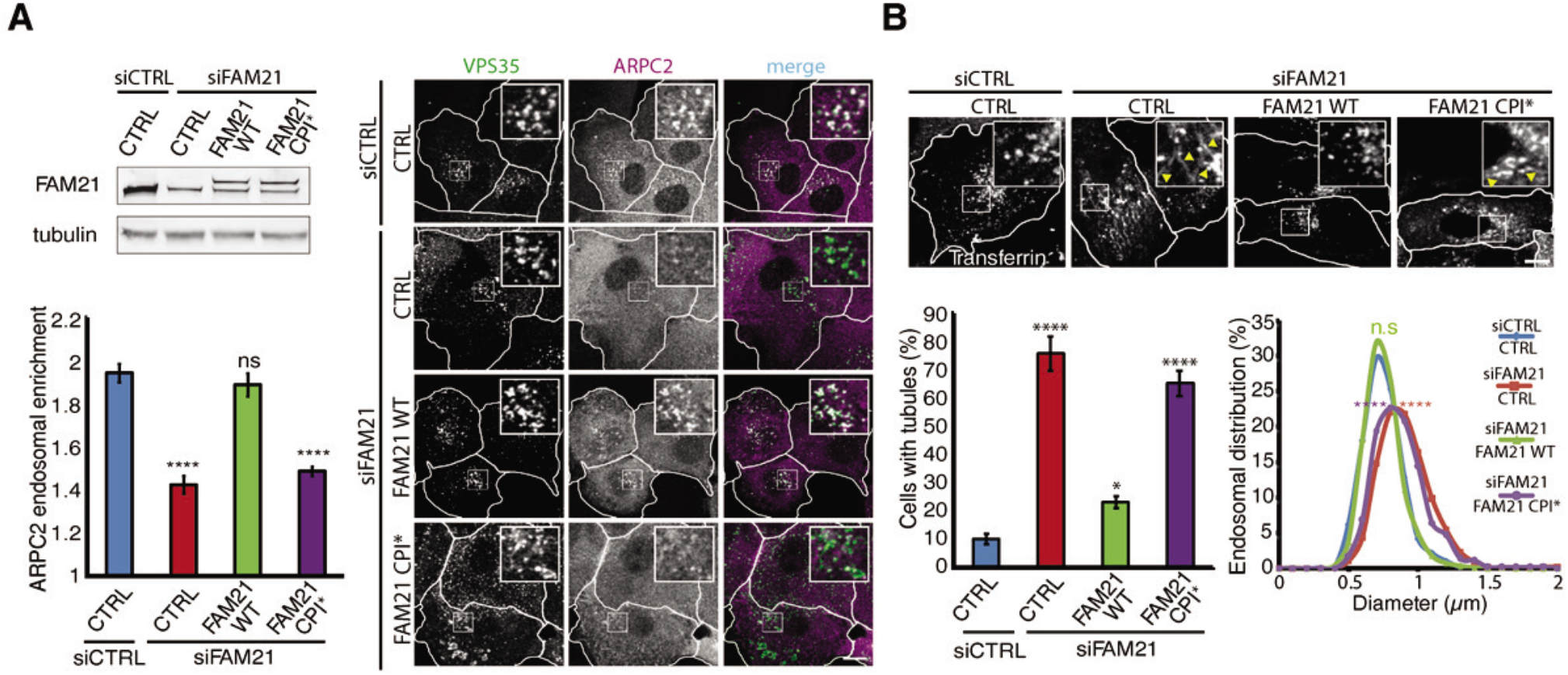
Deficient uncapping alters endosomal structures. **A** Endogenous FAM21 was depleted from MCF10A control cells or stable MCF10A lines expressing GFP-FAM21 WT or CPI* using siRNAs. Endosomal branched actin structures were estimated by the overlap of ARPC2 (Arp2/3) with VPS35 (retromer). Mean ± s.e.m., two-tailed Kruskal-Wallis test. **** P<0.0001. n=3. Scale bar: 10 μm. **B** Endosomes were loaded at steady state with fluorescent transferrin. Endosomal size and the presence of tubules (arrowheads) were estimated from live cell confocal imaging. Tubulation: Mean ± S.D., two-tailed one-way ANOVA. n=3. Apparent endosomal diameter: twotailed one-way ANOVA. n=2. * P<0.05, **** P<0.0001.

We studied the recycling of two well established WASH-dependent cargoes, integrins (*24*, *25*) and the GLUT1 glucose transporter (*26*, *27*). Levels of β1 integrins were down-regulated in FAM21 depleted cells, because of their deficient recycling to the plasma membrane following internalization in absence of functional WASH complex (Fig. 5, A and B). FAM21 depleted cells were able to protrude efficiently, but several protrusions often stretched cells in different directions, resulting in frequent shape changes, captured by aspect ratio volatility (Fig.5C, Movie S3, fig. S9A). Overall, FAM21 depleted cells explored a larger surface area than controls (Fig.5C; fig. S9, B and C), as indicated by their mean square displacement. FAM21 wild type, but not CPI*, rescued β1 integrin recycling defects and aberrant cell migration. Recycling of GLUT1 from endosomes to the plasma membrane requires the sorting nexin SNX27 and the WASH complex (*26*). FAM21 knock-down accumulated GLUT1 in SNX27 positive endosomes at the expense of the plasma membrane. FAM21 wild type, but not FAM21 CPI*, restored GLUT1 at the plasma membrane (Fig. 5D). The lack of GLUT1 at the plasma membrane translated into a profound defect in the uptake of 2-NitroBenzoxaDeoxyGlucose(2-NBDG), a fluorescent glucose analog (Fig. 5E, fig. S9D).

**Fig. 5.**
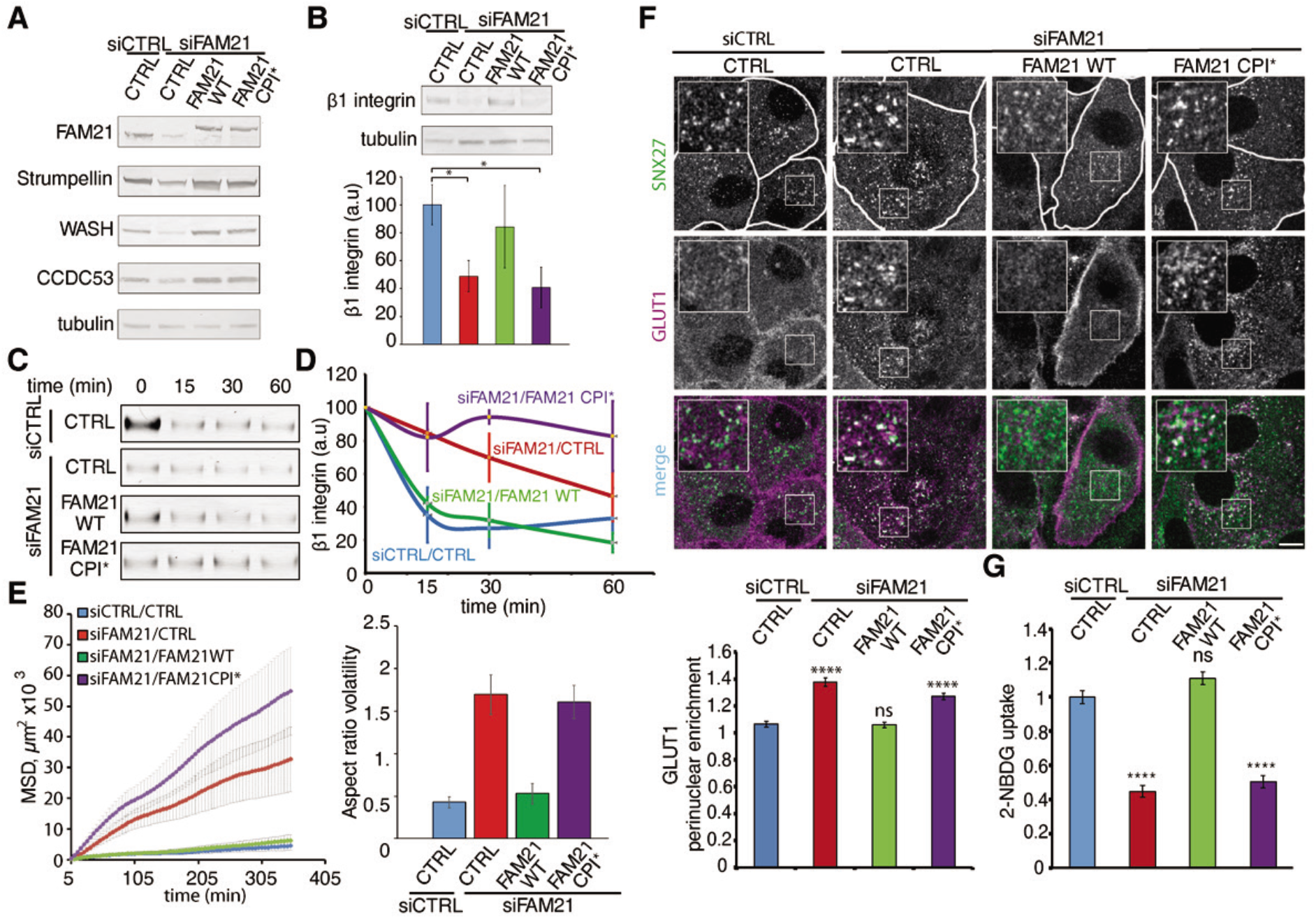
Uncapping by the WASH complex is required for cargo recycling. **A** Endogenous FAM21 was depleted from MCF10A control cells or stable MCF10A lines expressing GFP-FAM21 WT or CPI* using siRNAs. WASH complex subunits were analyzed by Western blots. **B** Levels of β1 integrin and quantification by densitometry. Mean ± s.e.m, paired t-test, n=3, * P<0.05. **C** β1 integrin recycling. Surface proteins that were biotinylated and internalized were allowed to recycle to the plasma membrane for the indicated time, before extracellular exposed biotin was removed. Streptavidin captured protein were analyzed by β1 integrin Western blots. **D** Quantification by densitometry. Areas under curve of siFAM21 and CPI*, but not of WT, are significantly different from siCTRL (P<0.05, ANOVA). n=3. **E** Mean Square Displacement (MSD) and aspect ratio volatility of single cells. ANOVA followed by Dunnett’s multiple comparisons, n=2. **F** The glucose transporter GLUT1 localizes to perinuclear SNX27-positive endosomes and at the plasma membrane. Scale bar: 10 μm. Mean ± s.e.m., two-tailed Kruskal-Wallis, n=3. **G** Cell uptake of the glucose analogue 2-NBDG. Mean ± s.e.m., two-tailed Kruskal-Wallis, n=2. **** P<0.0001.

The lack of rescue by FAM21 CPI* in these structural and functional analyses is striking considering that essential endosomal machinery is present in these experiments and that the only defect in the WASH complex reconstituted with FAM21 CPI* is its ability to interact with CP. Together, our in vitro reconstitutions and cell biology experiments support a working model where the WASH complex coordinates two multiprotein complexes containing distinct actin-related proteins (fig. S9E, Movie S4). The WASH complex is the only stable multiprotein complex that combines CPI and VCA motifs. WASH first uncaps Dynactin through the CPI of FAM21. The barbed end elongation from uncapped Dynactin then provides a primer actin filament for the Arp2/3 autocatalytic reaction.

We found that the CPI motifs of CARMIL and CIN85 can also induce actin elongation by uncapping Dynactin in vitro (Fig. S10). These two proteins, however, which associate with dendritic actin networks at the lamellipodial edge (*28*, *29*) and clathrin coated pits (*30*, *31*), respectively, do not appear to colocalize with Dynactin in the cell (Movies S5 and S6). Uncapping Dynactin to provide a primer filament to the Arp2/3 branching reaction is critical at the surface of endosomes, but probably not at the cell cortex, which is already rich in actin filaments.

Our newly discovered role of Dynactin in regulating actin polymerization complements its well-characterized function in promoting transport along microtubule tracks. Dynactin is organized around a central Arp minifilament, which recruits Dynein adaptor proteins along its length (*32*) and that primes the Arp2/3 complex via the elongation of an actin filament from its barbed end. Actin and microtubule cytoskeletons are both involved in controlling shape, motility and scission of endosomes. Dynactin thus plays a critical role towards both of these cytoskeletal elements at the surface of endosomes.

## Acknowledgments

We thank Pekka Lappalainen and Daniel Billadeau for kind gifts of DNA constructs, Laurent Blanchoin for the kind gift of Profilin, Pierre Mahou for assistance in confocal microscopy, Jérôme Boulanger for image processing advice.

## Funding

This work was supported in AG’s group by grants from the Agence Nationale de la Recherche (ANR ANR-15-CE13-0016-01), from the Fondation ARC pour la Recherche sur le Cancer (PGA120140200831), from Institut National du Cancer (INCA_6521). ED is supported by the Medical Research Council (MC_UP_1201/13) and the Human Frontier Science Program (CDA00034/2017). The imaging facility of Laboratoire d’Optique et Biosciences (Ecole Polytechnique) is partly supported by Agence Nationale de la Recherche (ANR-11-EQPX-0029 Morphoscope2).

## Author contributions

AF established most stable cell lines, characterized them and performed all cell biology experiments. VD purified most recombinant proteins and performed pyrene actin assays. CLC supervised some actin polymerization assays. NR purified some recombinant proteins and the dually tagged Dynactin by tandem affinity purification. LC and GRL performed in vitro TIRF assays. GRL supervised all of them. CES purified pig brain Dynactin and performed gel filtration analysis. CES and APC performed EM reconstruction of uncapped Dynactin. MAN performed ITC measurements. KO cloned Dynactin subunits and created the animated model. KO and MVH performed the coIP in brain extracts under the supervision of FS. ED performed live confocal microscopy. VH prepared several DNA constructs. NM constructed an expression vector and derived the corresponding stable cell line. AF drafted the manuscript. AMG supervised the work and wrote the manuscript. AF and VD contributed to the conceptual development of the project.

## Competing interests

The authors declare no competing financial interests.

## Data and materials availability

All data and the custom code to assess the 3 color colocalization are available from the authors upon reasonable request.

## Notes

### Competing Interest Statement

The authors have declared no competing interest.

