## Supplementary Materials for "The Arp1/11 Minifilament of Dynactin Primes the Endosomal Arp2/3 Complex"

**This PDF file includes:**

Materials and Methods
Figs. S1 to S10
Captions for Movies S1 to S6

**Other Supplementary Materials for this manuscript include the following:**

Movies S1 to S6

### Materials and Methods

Plasmids

All ORFs, or fragments thereof, were flanked by FseI and AscI restriction sites for easy shuttling between compatible plasmids. All PCR amplified fragments were sequenced to ensure that no unwanted mutation was introduced. Custom made plasmids were created using the MXS building block strategy (*33*).

WASH complex. FAM21 (FAM21C, NP_001317003) was amplified from the pCM3.H1p. shFAM21C.HA YFP FAM21C construct provided by Daniel Billadeau (Mayo Clinic, Rochester, MN, USA). FAM21 CPI corresponds to the fragment called U3A (*14*), i.e the peptide 934-1071 (ETPQ – QWAD). The CPI* mutations L1024A, R1031A, P1040A were introduced into the CPI fragment or full-length FAM21 by Quikchange site-directed mutagenesis (Agilent). WASH ORF and VCA were previously described (*2*).

Arp2/3 complex. ARPC5 and ARPC1B were amplified from described plasmids (*34, 35*).

Dynactin. ORFs encoding human Arp1, DCTN3 and DCTN6 were amplified by PCR from IMAGE clones number 3009822, 6454606 and 3925231, respectively.

CARMIL1 (also known as LRRC16A) was amplified from IMAGE clone 9021638. CARMIL1 CPI corresponds to the fragment called CBR (*29*), i.e. peptide 964-1078 (EKRS - KSRS).

CIN85 (also known as SH3KBP1) was amplified from IMAGE clone 3906722. CIN85 CPI corresponds to the peptide 426-523 (VGPL – ISLA).

CPα was amplified by PCR from pRFSDuet encoding the His-tagged α1 and β2 subunits of murine Capping Protein was from Pekka Lappalainen (University of Helsinki, Finland).

The GFP-binding protein (GBP) was obtained by gene synthesis (*36*).

Plasmids for protein production. CPI peptides from FAM21, CARMIL1 and CIN85 were cloned into pET28b for *E.coli* expression as a His tag fusion protein. FAM21 CPI and CPI* were also cloned into pGEX CS for *E.coli* expression as a GST fusion protein. pGEX-WASH VCA was described (*2*). pRFSDuet encoding the His-tagged α1 and β2 subunits of murine Capping Protein allows purification of the CP heterodimer. The GFP-binding protein (GBP) was cloned into a modified pET28b vector encoding a fusion protein with His tag and the Avitag sequence, which is biotinylated by BirA.

Plasmids for expression in human cells.

Rescue plasmids. FAM21 CPI WT and CPI* were cloned into MXS AAVS1L SA2A Puro bGHpA PGK rtTA3 SV40pA TRE EGFP Blue SV40pA AAVS1R and MXS PGK Blasti bGHpA EF1Flag iRFP Blue SV40pA.

Tandem Affinity Purification. Arp1 and DCTN3 were inserted into pCDNAm FRT Flag-GFP vectors (*2*). CPα was inserted into MXS PGK Puro bGHpA EF1α 6His PC Cherry CapZα SV40pA.

Live cell imaging experiments required construction of:
MXS AAVS1L SA2A Puro bGHpA EF1Flag EGFP ARPC5 SV40pA AAVS1R
MXS PGK Puro bGHpA EF1Flag EGFP CARMIL1 SV40pA
MXS PGK Puro bGHpA EF1Flag EGFP CIN85 SV40pA
MXS PGK Zeo SV40pA EF1Flag mCherry DCTN6 SV40pA
MXS PGK Zeo SV40pA EF1Flag mCherry Arp1 SV40pA
MXS PGK Blasti SV40pA EF1Flag iRFP WASH SV40pA
MXS PGK Blasti SV40pA EF1Flag iRFP ARPC1B SV40pA

<u>Cells and Transfections</u>

The immortalized epithelial cell line from human breast, MCF10A, was cultured in DMEM/F12 medium (Gibco) supplemented with 5% horse serum (Sigma), 100 ng/mL cholera toxin (Sigma), 20 ng/mL epidermal growth factor (Sigma), 0.01 mg/mL insulin (Sigma), 500 ng/mL hydrocortisone (Sigma) and 100 U/mL penicillin/streptomycin (Gibco). These cells were transfected with plasmids using Lipofectamine 3000 (Invitrogen). To obtain homologous recombination at the well-expressed AAVS1 locus, constructs were cotransfected with AAVS1 TALEN constructs (Addgene #59025 and 59026) (*37*). 2 days after transfection, the appropriate selective antibiotic is added (1 µg/mL Puromycin, 8 µg/mL Blasticidin, 100 µg/mL Zeocin). Single clones were isolated by cloning rings, expanded and characterized. To induce GFP-FAM21, two days of doxycycline treatment at 2 µg/mL were used. For siRNA-induced depletion of proteins, cells were transfected twice with 20 nM of targeting siRNA (Sigma) at day 0 and at day 2 using Lipofectamine RNAi max (Invitrogen). Cells were harvested or fixed at day 5. The following siRNAs were used:

CTRL - AAUUCUCCGAACGUGUCACGUUU,

FAM21 3′UTR - GCAAAUACUAGAACAGCUAGCUU,

ARP1 (ACTR1A) – GCUUUCUUGAGUCGGAGUGUUUU,

ARP11 – GCGCGUACUUGCUUUGUAAdTdT(*20*).

293 FlpIn TREx cells containing a single FRT site at a well-expressed locus (Invitrogen) were grown in DMEM supplemented with 10 % fetal calf serum (Invitrogen), 50 µg/mL zeocin (Invivogen) and 5 µg/mL blasticidin (Invivogen). Cells were transfected by calcium phosphate precipitation with pCDNAm plasmids expressing Flag-GFP tagged Dynactin subunits and the Flp

recombinase encoding pOG44 (Invitrogen). Pools of stable transfected cells were selected with 200 µg/mL hygromycin (Invivogen). These stable cells were then transfected with MXS PGK Puro bGHpA EF1Flag mCherry CPα SV40pA and selected with 1 µg/mL of puromycin (Invivogen). Stable dual clones were isolated and expanded in spinners for a total volume of 6 L as described (*38*).

Antibodies

For immunofluorescence the following antibodies were used: GLUT1 rabbit pAb and SNX27 mouse mAb from Abcam (ab15310 and ab77799 respectively); VPS35 (B-5) was from Santa Cruz Biotechnology

For Western blots the following antibodies were used: polyclonal antibody to Strumpellin (C-14) and monoclonal antibodies to Arp1/ACTR1 (α/β centractin, E-5), polyclonal Arp3 (#07–272) was from Cell Signalling; p62/DCTN4 (H-4) were from Santa Cruz Biotechnology; ArpC5 monoclonal antibody (clone 323H3) was from Synaptic Systems; tubulin mAb (clone DM1A) was from Sigma; $p150^{Glued}$/DCTN1 and β1-integrin mAbs were from BD Biosciences (BD610474 and BD610468, respectively); GFP monoclonal antibodies (13.1 and 7.1 clones) were from Roche; CPα and β mAbs (5B12.3 and 3F2.3, respectively) were from Developmental Studies Hybridoma Bank; Rabbit pAb to p27/DCTN6 (16948-1-AP) was from Proteintech; home-made polyclonal rabbit antibody CCDC53 was previously described (*2*, *39*).

For both applications, the following antibodies were used: home-made polyclonal rabbit antibodies targeting WASH, and FAM21 were previously described (*2*, *39*); ARPC2 rabbit polyclonal antibody was from Millipore (07-227).

Secondary goat anti-mouse and anti-rabbit antibodies conjugated with Alexa Fluor 488, 555 and 647 used for immunofluorescence were from Life Technologies.

Secondary goat anti-mouse and anti-rabbit antibodies conjugated with alkaline phosphatase used for Western blots were from Promega.

Immunoprecipitations and Western Blots

Each mouse brain was homogenized in the following buffer: 50 mM HEPES pH 7.7, 100 mM KCl, 1 mM $MgCl_2$, 1% Triton X-100, 320 mM sucrose supplemented with the cOmplete, EDTA free Protease Inhibitor Cocktail (Roche). The homogenate was purified by 2 subsequent centrifugations at 4°C (10 min at 1000 xg and 40 min at 12,000 xg) to remove nuclei and obtain a crude cytoplasmic fraction used for immunoprecipitation. MCF10A cells expressing GFP-FAM21 were lysed in (50 mM HEPES, 200 mM NaCl, 1 mM $CaCl_2$, 5 mM $MgCl_2$, 5% Glycerol, 1% Triton X-100, pH 7.4) supplemented with protease inhibitors. Cell extracts were incubated with 15 µL of GFP-trap beads or RFP-trap beads (Chromotek) for 2 h at 4 °C, washed extensively with lysis buffer and resuspended in SDS loading buffer. SDS-PAGE was performed using NuPAGE 4–12

% Bis-Tris or 3-8 % Tris-Acetate gels (ThermoFischer Scientific). After transfer, nitrocellulose membranes were blocked in 5 % milk and incubated with primary and secondary antibodies. For Western blots analyzed by densitometry, intensity of a rectangle area containing the band of interest with a subtracted background was divided by the intensity of a similar rectangle containing α-tubulin staining of the same blotting membrane. The intensities were normalized to 100 % in control conditions and plotted. For the densitometry analysis of Dynactin subunits in Fig.2B, values for each Dynactin subunit was normalized by the area under curve.

Sucrose Gradient

Nitrogen cavitation (Parr instruments, 500 Psi for 20 min) followed by centrifugation (16,000 × g, 20 min) and ultracentrifugation (150,000 × g, 60 min) were used to prepare cytosolic extracts from cells trypsinized from two 15 cm dishes and resuspended in the XB buffer (20 mM HEPES, 100mM NaCl, 1mM MgCl2, 0.1 mM EDTA, 1mM DTT, pH 7.7). 200 µL of extract was loaded on the 11 mL 5–20% sucrose gradient in the XB buffer and subjected to ultracentrifugation for 17 h at 197,000 ×g in the swinging bucket rotor SW41 Ti (Beckman). 0.5 mL fractions were collected and concentrated by using trichloroacetic acid precipitation with insulin as a carrier. The samples were washed with acetone, dried and then resuspended in the 1x LDS loading buffer with 2.5 % of βME for Western blot analysis.

Recombinant Protein Purification

All recombinant proteins were expressed in the BL21* strain (Invitrogen). The WASH VCA fragment was purified in fusion with GST as described (2). After concentration on Amicon Ultra (Millipore), it was dialyzed against (20 mM Tris pH 7.5, 25 mM KCl, 1 mM MgCl2, 150 µM sucrose), flash frozen in liquid nitrogen and kept at -80°C until use. GST-CPI and GST-CPI* were purified by glutathione fast-flow sepharose resin (GE Healthcare), followed by ion exchange chromatography. His-tagged CPI, CPI* and CP α1β2 were purified by immobilized metal ion affinity chromatography followed by anion exchange chromatography. NaCl-eluted fractions containing CPI and CP were dialyzed against (20 mM Tris pH 7.8, 40 mM KCl, 0.5 mM DTT, 1 mM EDTA, 20 % glycerol) for storage at -80°C. The GFP binding protein, His-Avi-GBP, was expressed and biotinylated during IPTG induction in the BL21* strain transformed with pACYC184-BirA. His-Avi-GBP was purified by Ni-NTA agarose affinity chromatography followed by anion exchange chromatography. NaCl-eluted fractions containing GBP were dialyzed against GBP buffer (20 mM Tris pH 8.0, 100 mM KCl, 2 mM DTT, 1 mM EDTA, 4 % glycerol) for storage at -80 °C. Purity and protein concentrations were determined after SDS-PAGE using purified actin as a standard, because it is accurately measured by its absorbance at 280 nM. Quantification by densitometry was performed with ImageJ.

Dynactin Purification

Native Dynactin was purified from pig brains using the previously described large-scale SP Sepharose protocol (*11*). Tagged Dynactin was purified from 293 stable cell lines expressing His-mCherry-CPα and Flag-GFP-Arp1 or Flag-GFP-DCTN3. 293 cell pellets were solubilized in (50 mM Tris pH 7.5, 150 mM NaCl, 1% Triton X-100, 4% glycerol, 5 mM $MgSO_4$, 0.1 mM $CaCl_2$, 0.1 mM ATP) supplemented with protease inhibitors. After clarification by centrifugation at 21,000 xg for 40 min, extracts were incubated with equilibrated anti-Flag M2 resin (Sigma). M2 beads were washed in (50 mM Tris pH 7.5, 150 mM NaCl, 0.1 % Triton X-100, 0.4 % glycerol, 1 mM $MgSO_4$, 0.1 mM $CaCl_2$), and eluted by 150 ng/µL 3xFlag peptide (Sigma) in (35 mM Tris pH 7.5, 150 mM KCl, 0.4 M sucrose, 1 mM $MgCl_2$, 0.1 mM $CaCl_2$, 0.2 mM ATP, 0.1 mM DTT) for 1 h at 18°C. Eluates were then incubated 3 h at 4°C with equilibrated Ni sepharose High Performance (GE Healthcare). Resin was washed in (35mM Tris pH 7.5, 150mM KCl, 0.5M sucrose, 1mM $MgCl_2$, 0.1mM $CaCl_2$, 0.1mM DTT, 0.2mM ATP, 5mM imidazole), and eluted in (35mM Tris pH 7.5, 150mM KCl, 0.5M sucrose, 1mM $MgCl_2$, 0.1mM $CaCl_2$, 0.1mM DTT, 0.2mM ATP, 150mM imidazole).

Gel Filtration Uncapping Assays

1.5 µM Dynactin was incubated for 5 min at room temperature alone with 3 µM FAM21 CPI, FAM21 CPI*, 15 µM CARMIL1 CPI, or not, in buffer (5 mM Tris pH 7.8, 100 mM KCl, 1 mM MgCl2, 0.1 mM CaCl2, 0.2 mM ATP (pH 7.0), 5 mM DTT). The mixture was then fractionated on a Superose 6 increase 3.2/300 column on an AKTAmicro system (GE Healthcare). The column was pre-equilibrated and run in the same buffer supplemented with the corresponding CPI.

Negative Stain EM

Dynactin was taken from the peak fraction and diluted 1:3 in buffer (5 mM Tris pH 7.8, 100 mM KCl, 1 mM MgCl2, 0.1 mM CaCl2, 0.2 mM ATP (pH 7.5-7.8), 5 mM DTT). Protein concentrations were chosen to give densely packed particles (approximately 400 per image).

400 mesh copper grids coated with a continuous carbon support layer (Agar scientific) were treated by glow discharging at 25 mA for 45 s (PELCO EasiGlow). 4 µl of sample was applied to the grid and incubated for 60 s before blotting with filter paper. The grid was then washed in 4x 20 µl 2 % uranyl acetate and blotted again before being allowed to dry. 500 micrographs per sample were manually collected on a FEI Spirit T12 microscope equipped with Gatan 2K × 2K CCD (model 984). Micrographs were taken with 0.5–2.5 µm underfocus, at a nominal magnification of 11,000x with the digital pixel size 4.86 Å.

Micrographs were CTF corrected using GCTF (*40*). All subsequent processing steps were carried out in RELION-3.0 (*41*). A small set of particles were manually picked, 2D classified, and the filament-like classes chosen as a template for AutoPicking. Particles were subjected to further 2D classification before 3D classification and 3D refinement using a negative stain structure of Dynactin (*42*) as a reference. The cryo-EM structure of Dynactin (PDB: 5ADX) (*11*) was fit into the 3D negative stain maps using UCSF Chimera (*43*).

Actin Polymerization Assays

Actin was purified from rabbit skeletal muscle acetone powder and kept at 4°C in G buffer (5 mM Tris-HCl, pH 7.8, 0.2 mM ATP, 0.1 mM $CaCl_2$, 0.1 mM DTT, 0.01% $NaN_3$). Actin was labeled on cysteine 374 with pyrene or labeled on lysines with Alexa-488 or Alexa-568 using standard procedures. The Arp2/3 complex was purchased from Cytoskeleton, Inc. Human profilin 1 was expressed and purified in *E. coli* using a polyproline affinity column, 8 M urea eluted proteins were dialyzed against profilin storage buffer (10 mM Tris-HCl, pH 7.5, 0.1 mM EDTA, 50 mM KCl, 1 mM DTT) and kept at -80°C. Actin and profilin concentrations were determined by absorbance at 280 nm.

Profilin-actin complexes were formed by incubating G-actin with a 6-8 fold molar excess of profilin for 10 min on ice before initiation of polymerization. Dynactin was added to the mix in G buffer. The pyrene actin assay starts upon the addition of one-twentieth volume of 20x KME (2 M KCl, 20 mM $MgCl_2$, 4 mM EGTA). For control curves, an equal amount of buffer without protein was always included. Actin polymerization was monitored by the increase in fluorescence of pyrene-actin (λexc 366 nm, λem 407 nm) at 20°C in a Cary Eclipse spectrofluorimeter (Varian) with a multicell holder. Handling of data and graph drawing were performed using Kaleidagraph v4.03 software (Synergy Software).

Single Filament and Single Molecule Assays

Open flow chambers were formed by mounting a cleaned glass coverslip on a glass slide with parallel strips of double-sided tape (Fig.1F). The surface was passivated by incubating a solution of BSA and biotinylated BSA in the chamber. After rinsing, the chamber was incubated with neutravidin, rinsed, then incubated with 0.5 μM biotin-labeled GFP binding protein in GBP buffer. After rinsing, 2 nM of Dynactin, or control buffer, was incubated in Dynactin buffer (20 mM Tris pH 7.7, 50 mM KCl, 1 mM $MgCl_2$, 0.1 mM $CaCl_2$, 10 mM DTT). After rinsing, a solution containing 1 μM G-actin (15 % Alexa488 labeled), 8 μM profilin, 30 nM CPI or not, and 0.14 % methylcellulose in F-buffer (5 mM Tris, pH 7.8, 0.2 mM ATP, 0. 1 mM $CaCl_2$, 10 mM DTT, 1 mM $MgCl_2$, 0.2 mM EGTA, 100 mM KCl, 1 mM 1,4-diazabicyclo[2.2.2]octane) was flown in and images were acquired in TIRF on an inverted microscope (Nikon TiE, with a 60x TIRF objective) using 100 mW tunable lasers (ILAS2, Gataca Systems), an EMCCD Evolve camera (Photometrics) and Metamorph software.

The microfluidics experiment (Fig.1G) was performed in chambers made of Poly Dimethyl Siloxane (Sylgard) and solution flows were controlled with a MFCS system (Fluigent) as previously described (*44*). Surfaces were passivated with PEG-Silane (overnight incubation), then exposed to PLL-PEG-biotin for 1 h, and to PLL-PEG for 10 min, then rinsed extensively. Surfaces were then exposed sequentially to neutravidin, biotin labeled GFP binding protein and Dynactin. Finally, a solution containing profilin, Alexa568-actin (15 % labeled) and CPI was flowed in, and images were acquired.

CP-Dynactin colocalization experiments (Fig.2A) were carried out in open flow chambers, passivated with BSA. Solutions of dynactin, pre-incubated with CPI or CPI* or only buffer, were flowed in, and the chamber was rinsed with buffer after 30 seconds.

For branching experiments (Fig.3A), preformed red filaments of Alexa568-actin (15 % labeled) obtained by a 12 min preincubation of 2 nM Dynactin with 270 nM CPI, 1 μM profilin and 0.8 μM Alexa568-actin (15 % labeled) in F-buffer, were then diluted 20-fold in a solution containing 60 nM WASH VCA 60 nM Arp2/3, 0.4 μM profilin and 0.4 μM Alexa 488-actin (15 % labeled, green) in F-buffer supplemented with 0.13% methylcellulose.The reaction was then flowed into a BSA passivated chamber.

Isothermal Titration Calorimetry

CP and CPI proteins were dialyzed against the ITC buffer (20 mM Tris, pH 7.5, 150 mM NaCl, 4% glycerol and 2 mM βME). Their interaction was analyzed using a MicroCal iTC200 microcalorimeter (GE Healthcare). Measurements were carried out in triplicate at 22 °C. GST-CPI, GST-CPI* or GST at 120 μM were stepwise injected (20 injections of 2 μl every 180 sec) from the syringe into the measurement cell containing 9 μM His-tagged CP. No interaction was measured between CP and GST. The change in heating power was observed over the reaction time until equilibrium was reached. Data were analyzed using the software provided by the manufacturer. The graphs show integrated heats of injection with the best fit to a one-site binding model using Origin 7.0 software.

Cell Treatments

For Transferrin uptake, cells were starved in DMEM supplemented with 0.5% of BSA and 10 mM HEPES pH7.4 for 30 min, then 10 μg/mL Transferrin coupled with Alexa 555 (ThermoFisher Scientific) was added for 30 min. Cells were then washed with starvation medium, before examination with the confocal microscope.

For 2-NBDG uptake, cells were extensively washed with glucose-free DMEM (Gibco), then 100 μg/mL 2-NBDG (N13195, ThermoFischer Scientific) was incubated for 10 min. Cells were then washed with glucose-free DMEM before examination with the confocal microscope.

For the endosomal branched actin regrowth assay, 1 μM LatrunculinA (Calbiochem) was incubated for 2 h, before being extensively washed away.

Recycling Assay

Cells in 6-well plates were washed with cold PBS. Cell surface proteins were labeled with 0.5 mg/mL of EZ-link cleavable sulfo-NHS-SS-biotin (Thermo Scientific) in Hank's balanced salt solution for 30 min at 4°C. Unbound biotin was washed away with cold medium, and pre-warmed serum containing medium was added to cells. Biotin-labeled surface proteins were allowed to internalize for 30 min at 37°C. Remaining surface exposed biotin was removed with 2 washes of reducing buffer (50 mM Tris–HCl, 100 mM NaCl, 60 mM MesNa, pH 8.6) for 15 min at 4°C, followed by quenching with (50 mM Tris-HCl, 100 mM NaCl, 100 mM iodoacetamide, pH 8.0) for 15 min on ice. The cells were incubated at 37°C for 0, 15, 30, 60 min and then exposed to reduction and quenching as above. Cells were washed with cold PBS and lysed by scraping in 100 µL of lysis buffer (50 mM Tris-HCl, 150 mM NaCl, 1.5% octylglucoside, 1% NP-40, 1 mM EDTA, pH 7.4 supplemented with protease inhibitors) and incubated at 4°C for 20 min. Cell extracts were cleared by centrifugation (16 000 × g, 20 min, 4°C). After BCA normalization, samples were incubated for 3 h with Streptavidin beads (ThermoFischer Scientific), washed 5 times and analyzed by Western blot.

Immunofluorescence

Cells were fixed either in 3.2% PFA prepared in PBS or in 1% PFA in serum-free DMEM followed by absolute methanol at -20°C in the case of GLUT1 staining. Cells were quenched with 50 mM $NH_4Cl$, then permeabilised with 0.5 % Triton X-100, blocked in 2 % BSA and were incubated with antibodies (1-5 µg/mL for the primary, 5 µg/mL for the secondary). Images were acquired by Leica SP8ST-WS confocal microscope equipped with a HC PL APO 63x/1.40 oil immersion objective, a white light laser, HyD and PMT detectors.

Live Fast Acquisition Microscopy

Live cells were plated on fibronectin-coated glass-bottom Petri dishes (World precision Instrument) for 2 h at 37 °C and imaged in Leibovitz's L-15 Medium without phenol red (Thermo Fisher) on a custom spinning disk confocal microscope composed of a Nikon Ti stand equipped with perfect focus, a 100X NA 1.49 SR TIRF objective, a Yokogawa CSU-X1 spinning disk head and a Photometrics 95B back-illuminated sCMOS camera operating in global shutter mode. Excitation was performed using 488 (150 mW OBIS LX), 561 (100 mW OBIS LS) and 637 nm (140 mW OBIS LX) lasers fibered by a Cairn laser launch. To minimize bleedthrough, single band emission filters were used (Chroma 525/50 for GFP, 595/50 for mCherry and 680/40 for iRFP) and acquisition of each channel was performed sequentially using a fast filter wheel (Cairn Optospin). Sample temperature was maintained at 37 °C using a temperature control chamber (Digital Pixel Microscopy System). Acquisition was controlled by Metamorph software.

For the TIRF imaging cells were spread overnight on fibronectin-coated glass (20 µg/mL in PBS 1 h for the coating) and imaged in Leibovitz's L-15 medium (Gibco) supplemented with 20 mM HEPES (Gibco) at 37°C using a temperature control chamber (Digital Pixel Microscopy System). TIRF imaging was performed on a custom-built TIRF system based a Nikon Ti stand equipped with perfect focus system, a fast Z piezo stage (ASI), a PLAN NA 1.45 100X objective and an azimuthal TIRF illuminator (iLas2, Roper France) modified to have an extended field of view (Cairn). Images were recorded with a Photometrics Prime 95B back-illuminated sCMOS camera run in pseudo-global shutter mode and synchronized with the azimuthal illumination. GFP (respectively mCherry, iRFP670) was excited by a 488 nm laser (respectively 561 and 637 nm, all Coherent OBIS mounted in a Cairn laser launch) and imaged using dedicated single bandpass filters for each channel mounted on a Cairn Optospin wheel (Chroma 525/50 for GFP and Chroma 595/50 for mCherry and 655LP for iRFP). TIRF angle was set independently for all channels so that the depth of the TIRF field was identical for all channels. System was operated by Metamorph.

To enhance the detection of dim endosomes in live imaging data (Fig. 3B, fig. S7 and Movie S2), a wavelet "à trous" filter (*9*) was applied to raw images and the resulting filtered image was averaged with the original raw image. This treatment was not applied when colocalisation was quantified. Endosome trajectories were used as the lines for kymograph measurement.

Videomicroscopy of Cell Migration

Videomicroscopy was performed on an inverted Axio Observer microscope (Zeiss) equipped with a Pecon Zeiss incubator XL multi S1 RED LS (Heating Unit XL S, Temp module, $CO_2$ module, Heating Insert PS and $CO_2$ cover), a definite focus module and a Hamamatsu camera C10600 Orca-R2. Pictures were taken every 5 min for 24 h using the Plan-Apochromat 20X/0.80 air objective. In the single cell migration assay, only the cells that were freely migrating for 12 h were taken into account. Cell trajectories were acquired with the ImageJ Manual tracking plugin. Analysis of cell migration was performed in the DiPer software (*45*). To measure aspect ratio, the ratio of the longest axis of a cell to the shortest one, cell boundaries were manually drawn. Volatility corresponds to its standard deviation at all time points.

Image Analysis

Image analysis was performed in ImageJ or FIJI software. To measure endosomal enrichment of ARPC2 staining, images were thresholded in the VPS35 channel by the Otsu algorithm to create a region of interest (ROI), then the mean intensity of the ARPC2 staining was measured in this ROI. Endosomal enrichment was defined as the ratio of mean ARPC2 intensity in the ROI divided by the mean intensity of ARPC2 in the whole cell. Similarly, perinuclear enrichment of GLUT1 was defined as the ratio of GLUT1 intensity in a disc of 100 $\mu m^2$ that contains most of GLUT1 signal in the perinuclear region divided by the one of the whole cell. Mean fluorescent intensities of 2-NBDG in the nucleus and in the whole cell were quantified. Cytoplasmic accumulation is defined as the difference between total and nuclear intensity.

To automatically measure colocalization between mCherry-DCTN6, GFP-ARPC5B and iRFP670-WASH, we used an object-based method, where two objects are considered delocalized if the distance between their fluorescence centroid ($d$) is below a certain threshold $d_{ref}$, usually set close to lateral resolution of the microscope ($r_{xy}$) (*46*). To segment endosomes, we used a threshold-free method based on 2D Gaussian fitting, which does not rely on an intensity threshold, but rather on the fact that particles have a Gaussian shape against the local background (Thunderstorm algorithm (*47*)). Once this automated segmentation has been performed in all 3 channels, the distance d between all objects in the iRFP670-WASH and mCherry-DCTN6 (respectively, GFP-ARPC5B) channels are computed and compared to the distance threshold ($d_{ref} = 0.36$ µm); measured using 0.2 µm TetraSpeck beads from Invitrogen). Once all the particles have been detected and their colocalization state addressed (i.e. $d < d_{ref}$), we measured the percentage of colocalisation. This measurement was averaged over five timepoints to avoid false positive colocalisation events due to particles detected in both channels at the same time by chance, then this was averaged between cells.

Counts and Statistics

Fig.3C: Number of cells/fields analyzed in each condition: siCTRL –61/19, 74/24, 79/23; siARP11 – 62/18, 101/32, 93/26; siARP1 – 74/27, 78/24, 38/30 (LatA, 3 min, 10 min).
Fig.4A: Number of cells/fields analyzed in each condition: siCTRL – 62/24; siFAM21 – 56/29; WT – 34/27; CPI* - 53/30.
Fig.4B: Tubulation: total number of cells: siCTRL – 127; siFAM21 – 106; WT – 107; CPI* – 125. Diameter: siCTRL – 1112/26; siFAM21 – 990/21; WT – 988/30; CPI* – 750/23.
Fig.5F: Number of cells analyzed: siCTRL – 42; siFAM21 – 68; WT – 46; CPI* – 71.
Fig.5G: Nnumber of cells analyzed: siCTRL – 110; siFAM21 – 110; WT – 101; CPI* – 102.

Statistical analysis was carried out with GraphPad Prism software (v7.00) and Microsoft Excel 2016. When not stated otherwise, ANOVA and Kruskal-Wallis test were used for the statistical analysis. Shapiro-Wilk normality test was applied to examine whether data samples fit a normal distribution. If data satisfied the normality criterion, ANOVA followed by post-hoc Tukey's multiple comparison test was performed. If not, non-parametric Kruskal-Wallis test followed by post-hoc Dunn's multiple comparison test was applied. 4 levels of significance were distinguished: * $p<0.05$, ** $p<0.01$, *** $p<0.001$, **** $p<0.0001$. n refers to the number of independent experiments.

**Supplementary Figures**

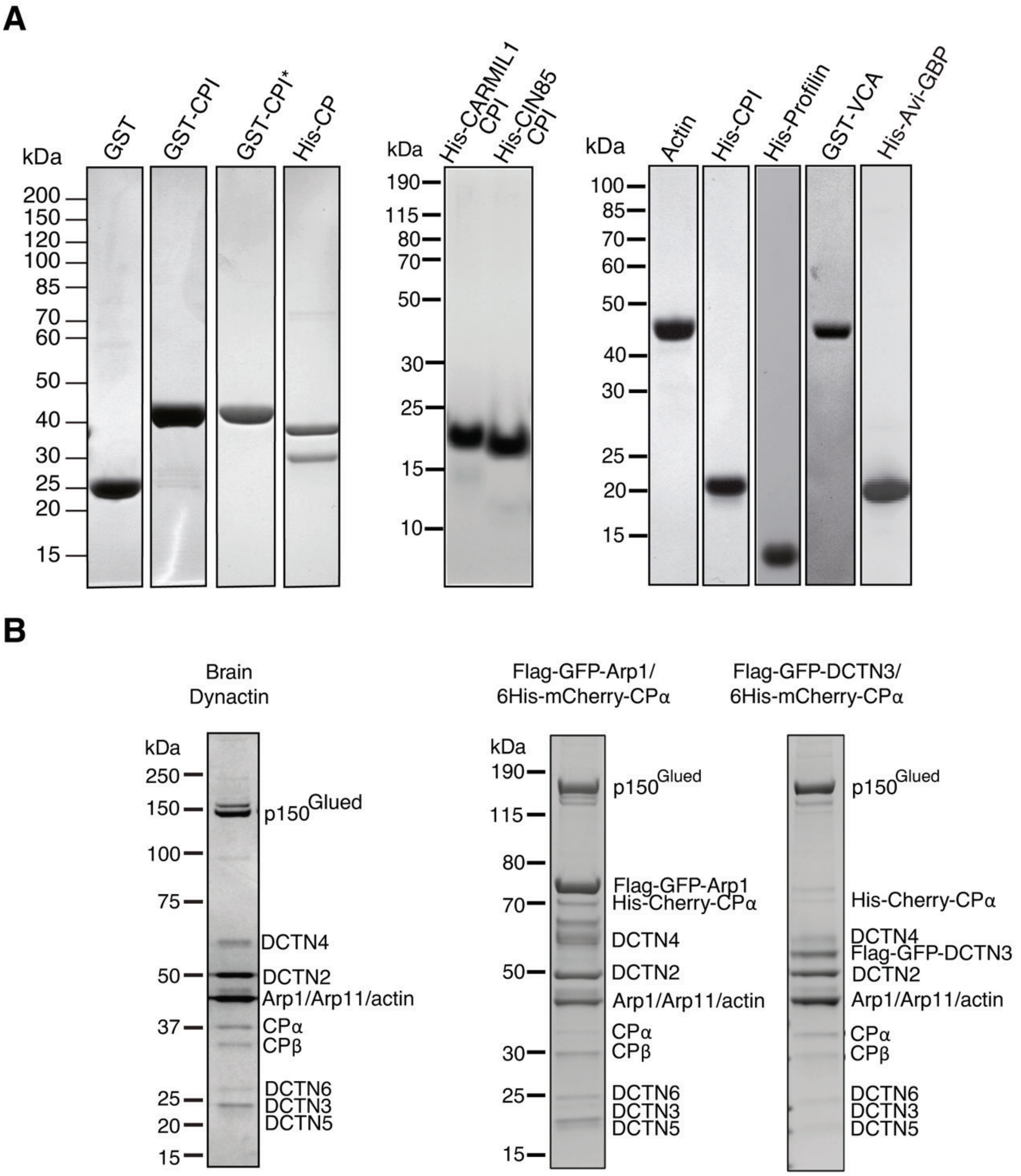


**Fig. S1. Purified proteins and complexes used in the study. A** SDS-PAGE of proteins expressed and purified from *E. coli*. His-Avi-GBP refers to the biotinylated GFP Binding Protein. **B** SDS-PAGE of purified Dynactins, native Dynactin from pig brain and labeled Dynactins purified from 293 stable cell lines by tandem affinity purification. Coomassie staining.

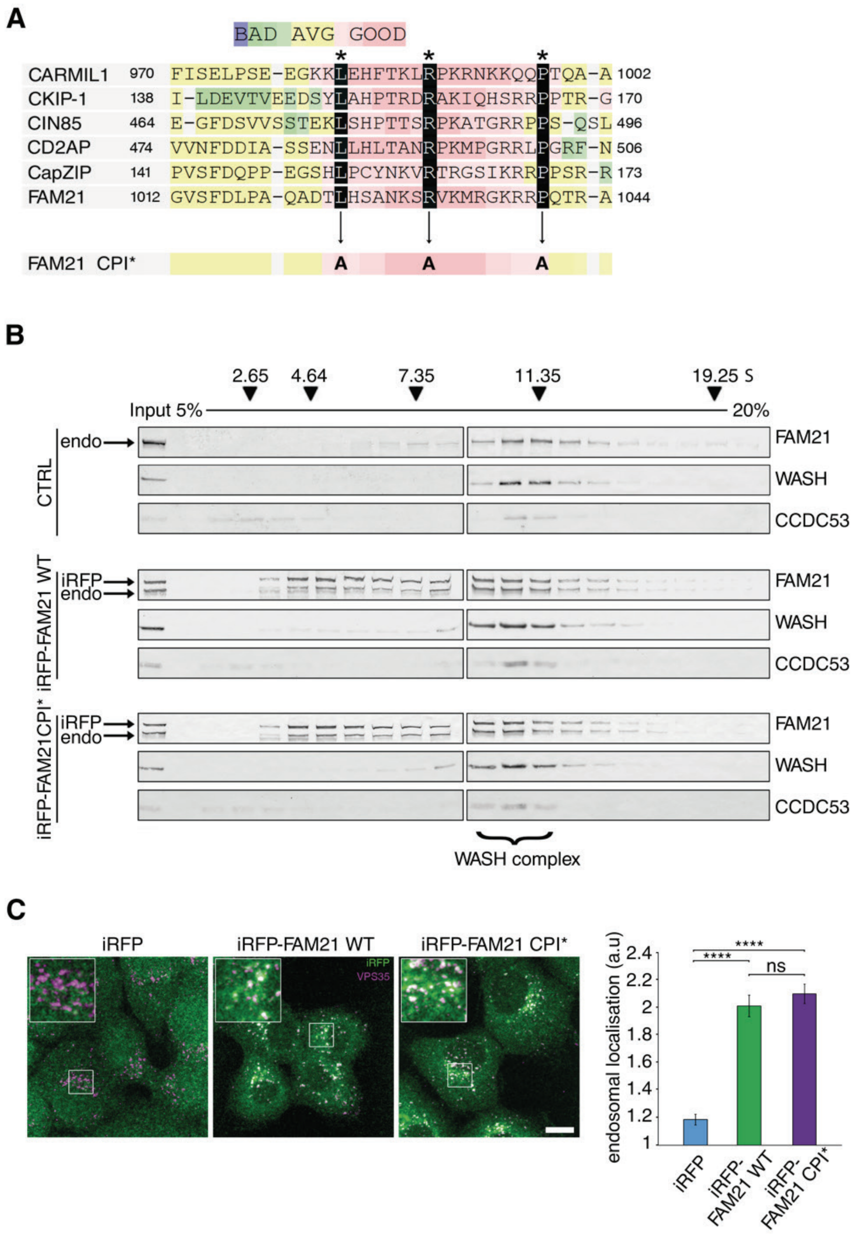


**Fig. S2. Characterization of FAM21 CPI*. A** Alignment of CPI motifs from CPI containing proteins(*6*) and mutations introduced to create CPI*. **B** Cytosolic extracts from parental MCF10A cells or stable clones expressing iRFP tagged FAM21 wild type or CPI* were fractionated by ultracentrifugation in sucrose gradients. Western blots of WASH complex subunits. WASH is only present within its 11 S complex. FAM21 wild type and CPI* are incorporated into the WASH complex, but also exist as lighter forms when overexpressed. Endogenous CCDC53 exists in a complexed form and in lighter forms. **C** MCF10A cell lines constitutively expressing iRFP, iRFP-FAM21 wild type or CPI* (green) were stained with VPS35 antibodies (magenta). Quantification of the iRFP signal in the endosomal object defined by VPS35 staining over the iRFP signal in the whole cell (43 iRFP cells, 55 iRFP-FAM21 cells and 52 iRFP-FAM21 CPI* cells analyzed). Scale bar: 10 µm. Mean ± s.e.m., two-tailed Kruskal-Wallis test, **** P<0.0001, ns: not significant.

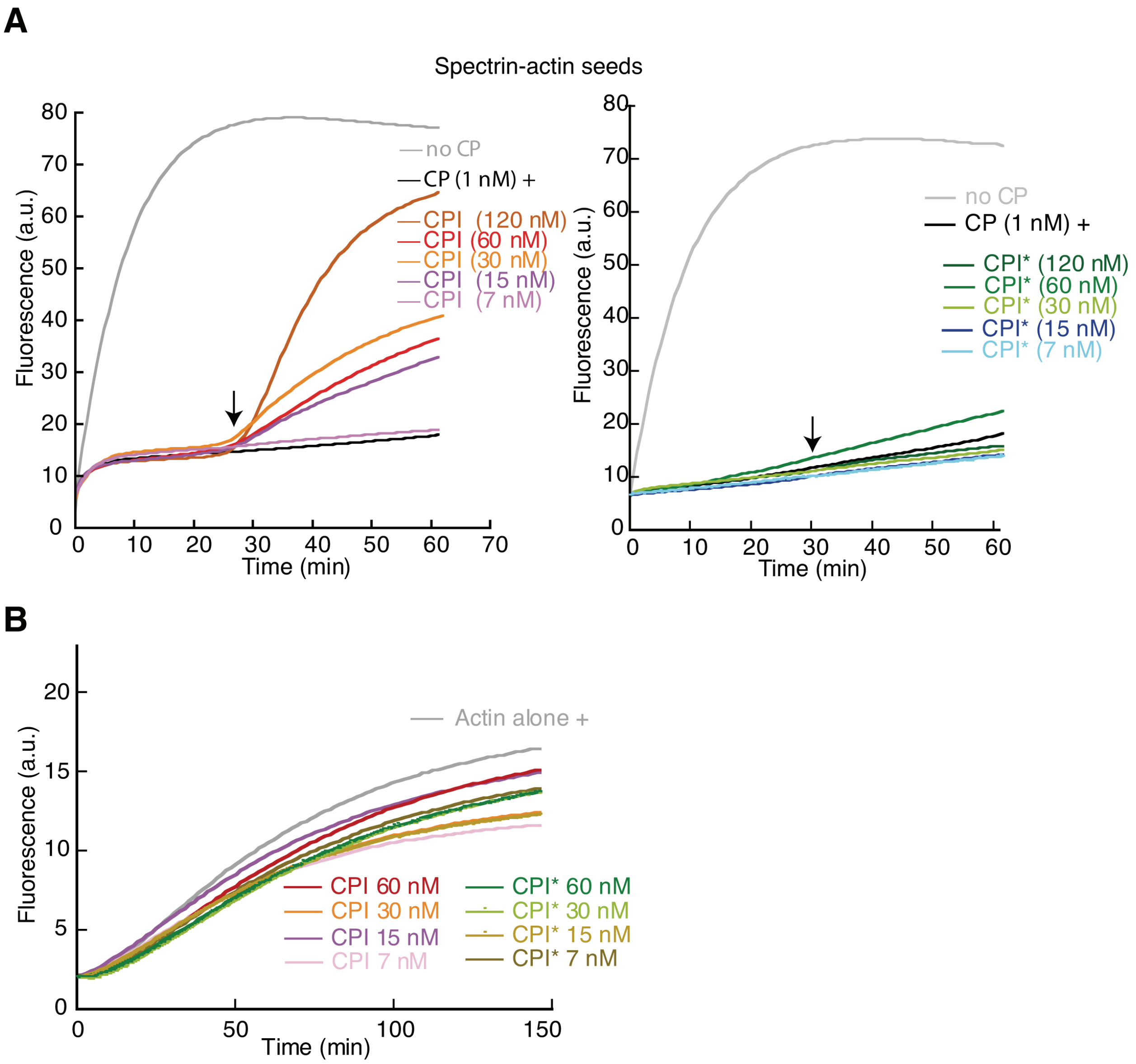


**Fig. S3. Comparison of CPI and CPI* in actin polymerization assays.** **A** Elongation of actin filaments from spectrin-actin seeds (120 pM) was interrupted or not by the addition of 1 nM CP. CPI, but not CPI*, uncaps actin filaments when added (arrow). **B** CPI and CPI* do not have a significant impact on actin polymerization in the absence of CP. Conditions: Pyrene-labeled actin (1.5 µM, 7% pyrene labeled). One representative experiment out of 3 is displayed.

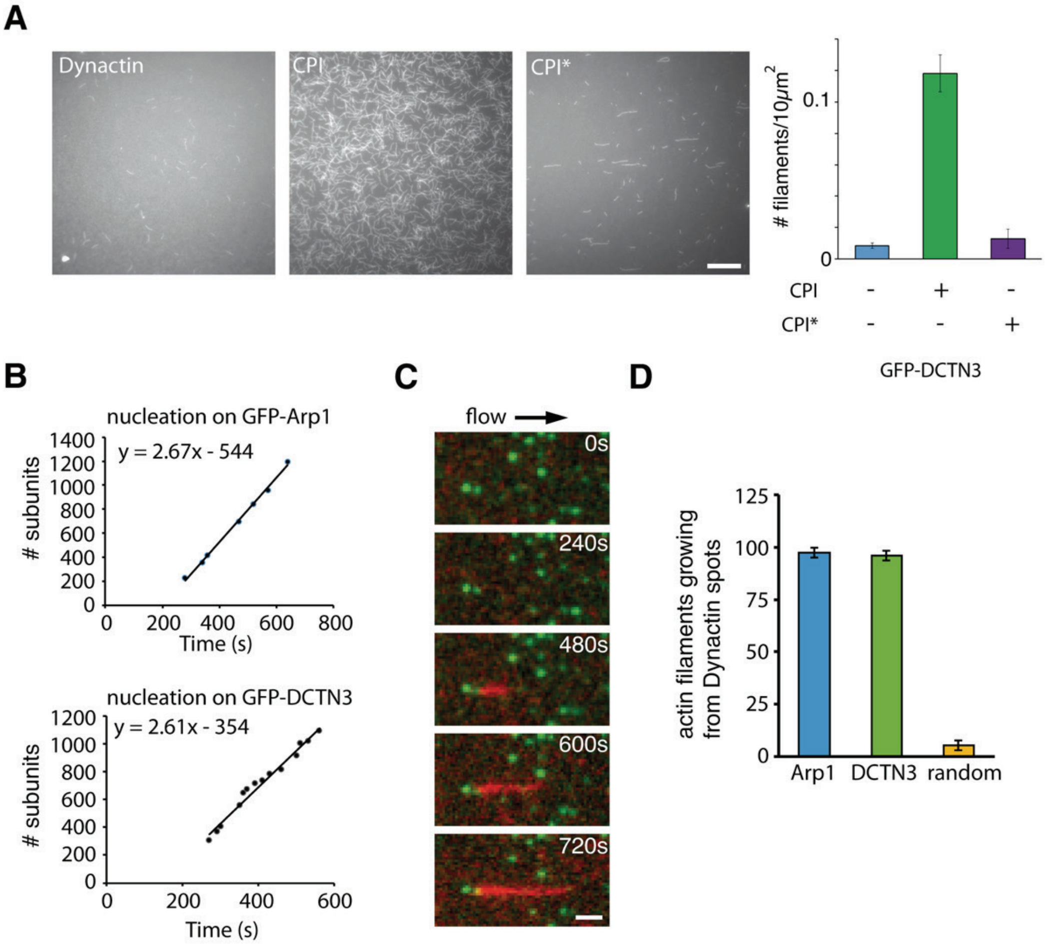


**Fig. S4. Growth of actin filaments from anchored Dynactin observed by TIRF microscopy.** **A** Nucleation of actin filaments in open chambers decorated with Dynactin tagged with GFP-DCTN3. Filament formation was observed 10-12 minutes after a solution of profilin-actin (1 µM 15% Alexa labeled actin, 8.2 µM profilin), with CPI or CPI* (2.7 µM) or without, in F buffer was flowed in. n=3 with two different preparations of Dynactin. **B** Rate of actin filament elongation (2 examples). Since each actin molecule contributes 2.7 nm of filament length, the measured elongation rate corresponds to 2-3 subunits per seconds, which is the characteristic rate of filament elongation from the barbed end with these concentrations of actin and profilin (*48*, *49*). **C** Time-lapse images of a filament growing under flow from anchored Dynactin. Conditions: 1 µM actin (15% Alexa488-labeled, red), 1 µM Profilin, 50 nM CPI, with a surface decorated with Dynactin (GFP-Arp1, green). Scale bar: 3 µm. **D** Quantification of actin filaments elongating from Dynactin spots. Random corresponds to images taken from different fields of view for each wavelength. n=3 for each type of tagged Dynactin, with two different preparations for GFP-Arp1 and for GFP-DCTN3, mean ± S.D.

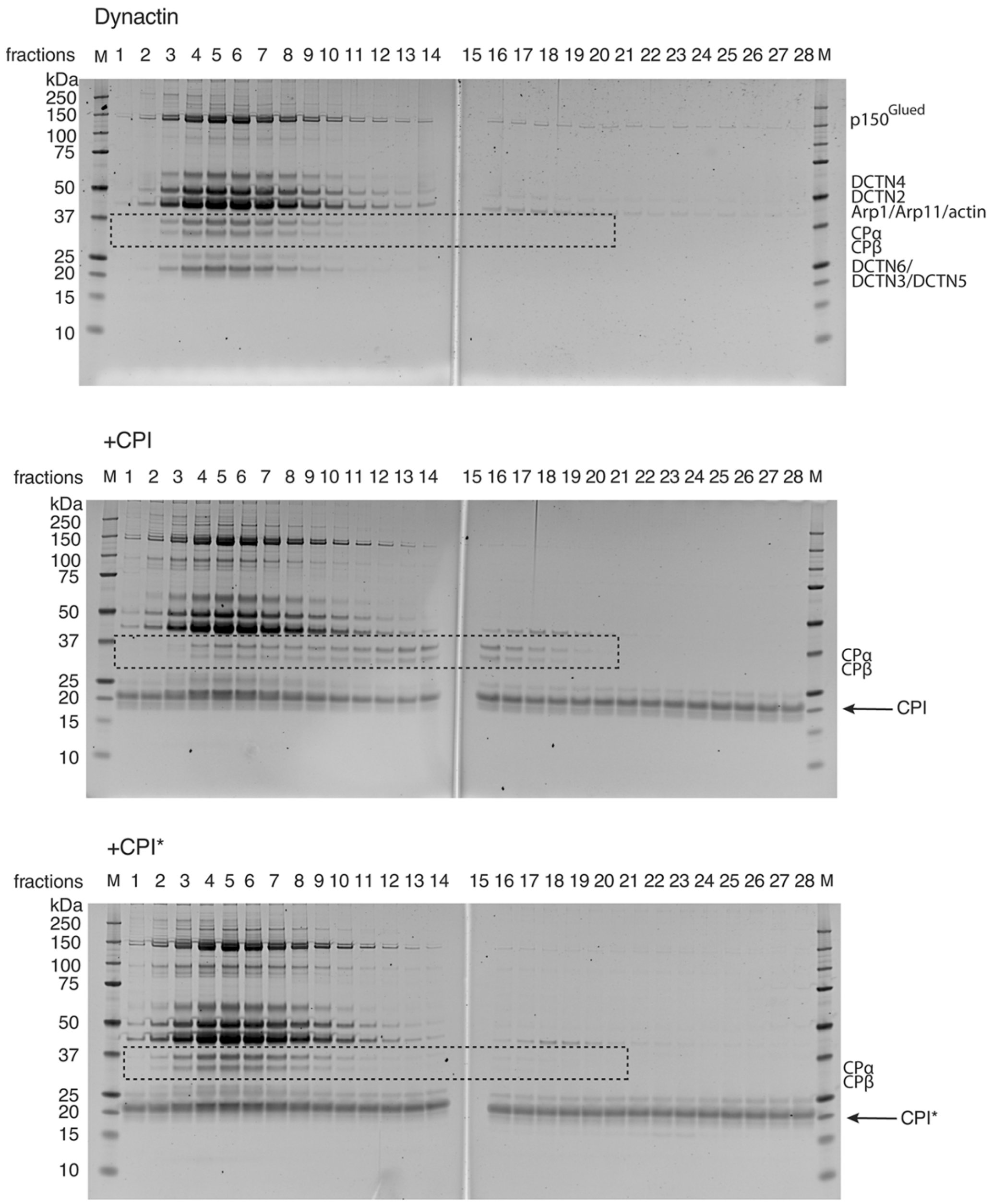


**Fig. S5. CPI induces the dissociation of CP from Dynactin.** Gel filtration of Dynactin (1.5 μM) in the continuous presence of CPI or CPI* (3 μM). Total proteins in fractions were stained with SYPRO Ruby. Dashed boxes highlight the CP that dissociates from Dynactin in the presence of CPI. n=3 independent experiments.

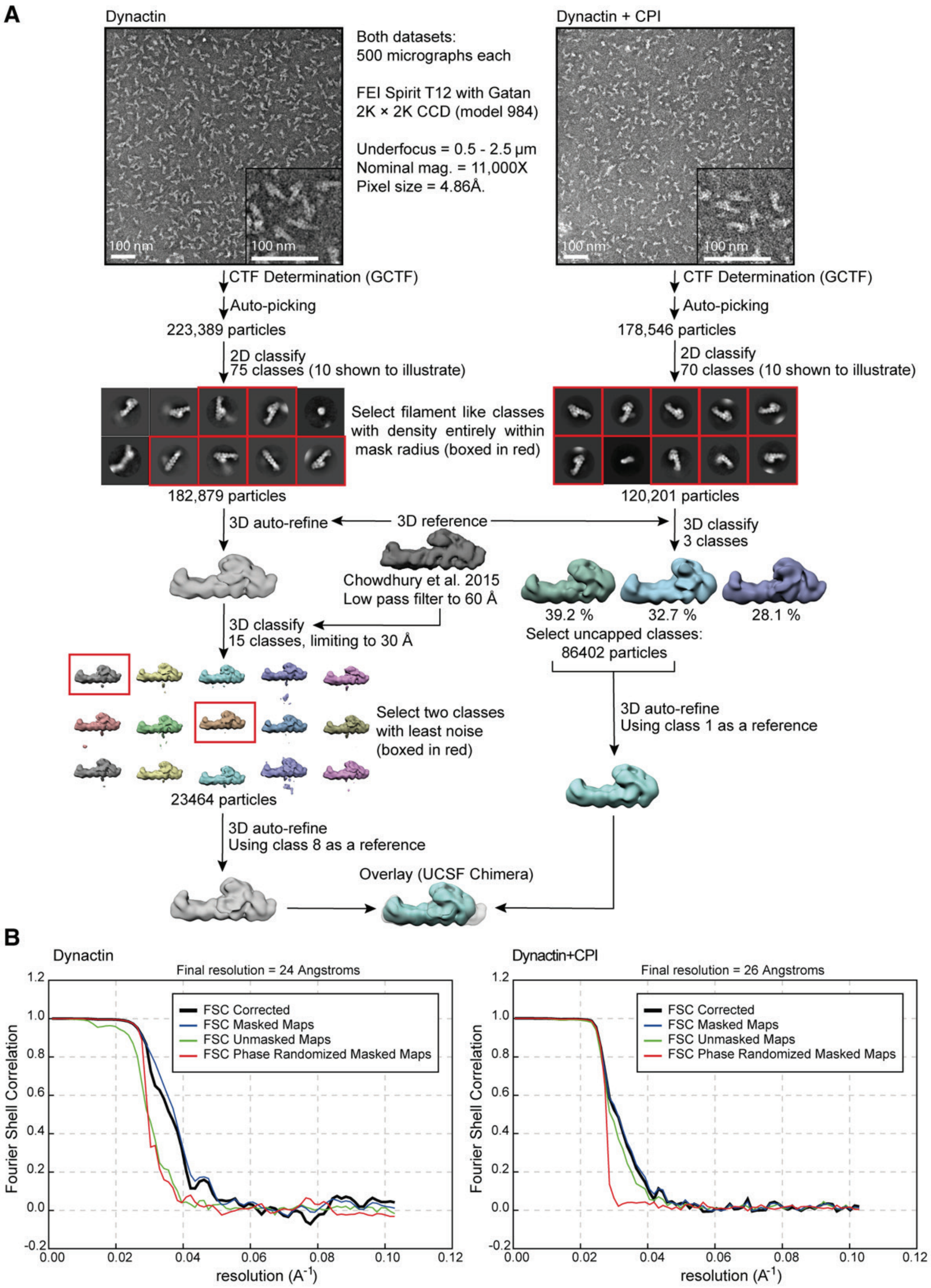


**Fig. S6. Negative Stain Electron Microscopy processing. A** Negative stain EM Relion processing pipeline for control Dynactin and Dynactin uncapped by CPI. 3D reference for Dynactin refinement was taken from reference (*42*). **B** Fourier Shell Correlation curves for control Dynactin and for Dynactin uncapped by CPI.

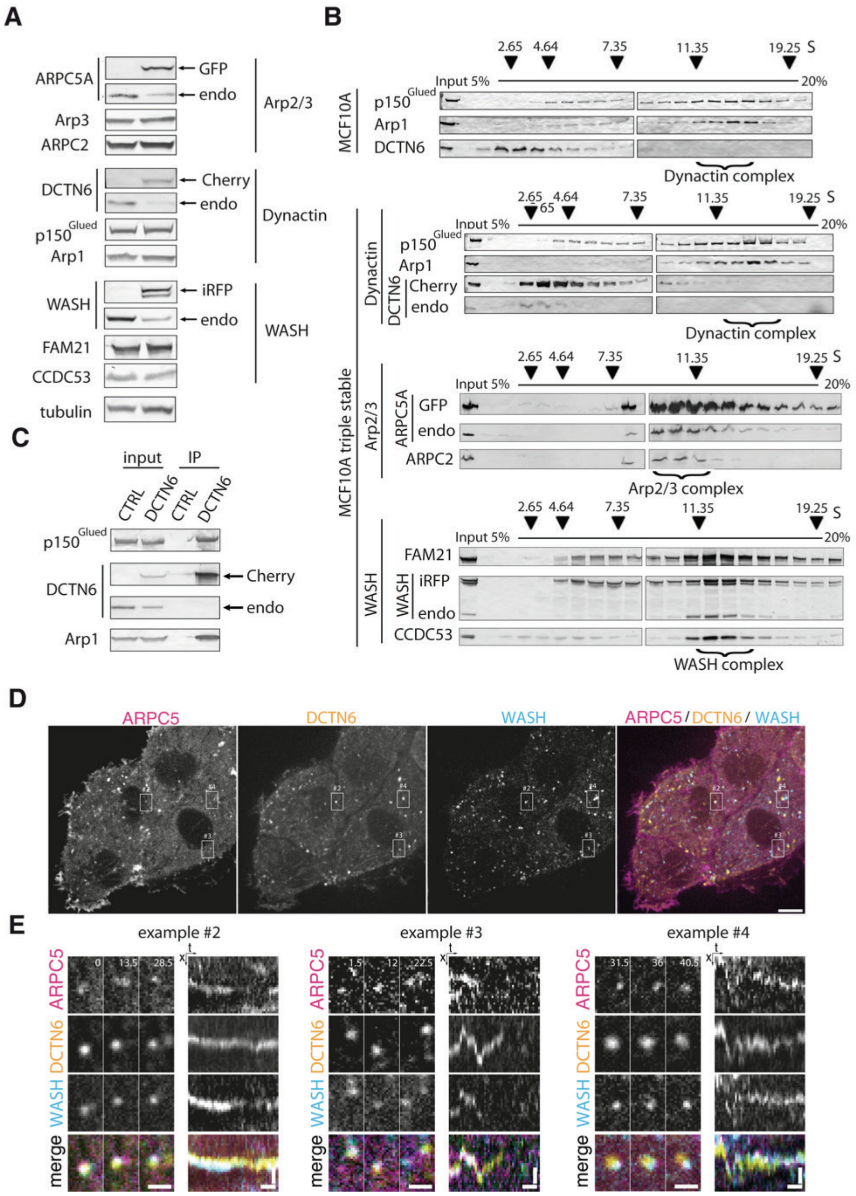


**Fig. S7. Characterization of the triple stable cell line expressing GFP-ARPC5, mCherry-DCTN6 and iRFP-WASH. A** Western blots of the triple transgenic line (right lane) compared to the parental MCF10A cell line (left lane). Tagged proteins replace their endogenous counterparts. **B** Cytosolic extracts were analyzed by ultracentrifugation on sucrose gradient. iRFP-WASH and GFP-ARPC5 were found in their respective complex. mCherry-DCTN6 and endogenous DCTN6 dissociate from Dynactin in sucrose gradients (*50*). **C** mCherry immunoprecipitations indicate that DCTN6 is incorporated into Dynactin. **D** Live spinning disk confocal microscopy (single plane). Same cell as in Fig.3B, with split channels to address colocalization. Scale bar: 6 µm. **E** Additional examples of colocalization (white boxes in D). Left panels show zoomed still images extracted from the movie, right panels associated kymographes along the trajectory of the object. Elapsed time in seconds. Scale bars: scale bar: 1.2 µm (left panels), 1.2 µm/12 s (right panels).

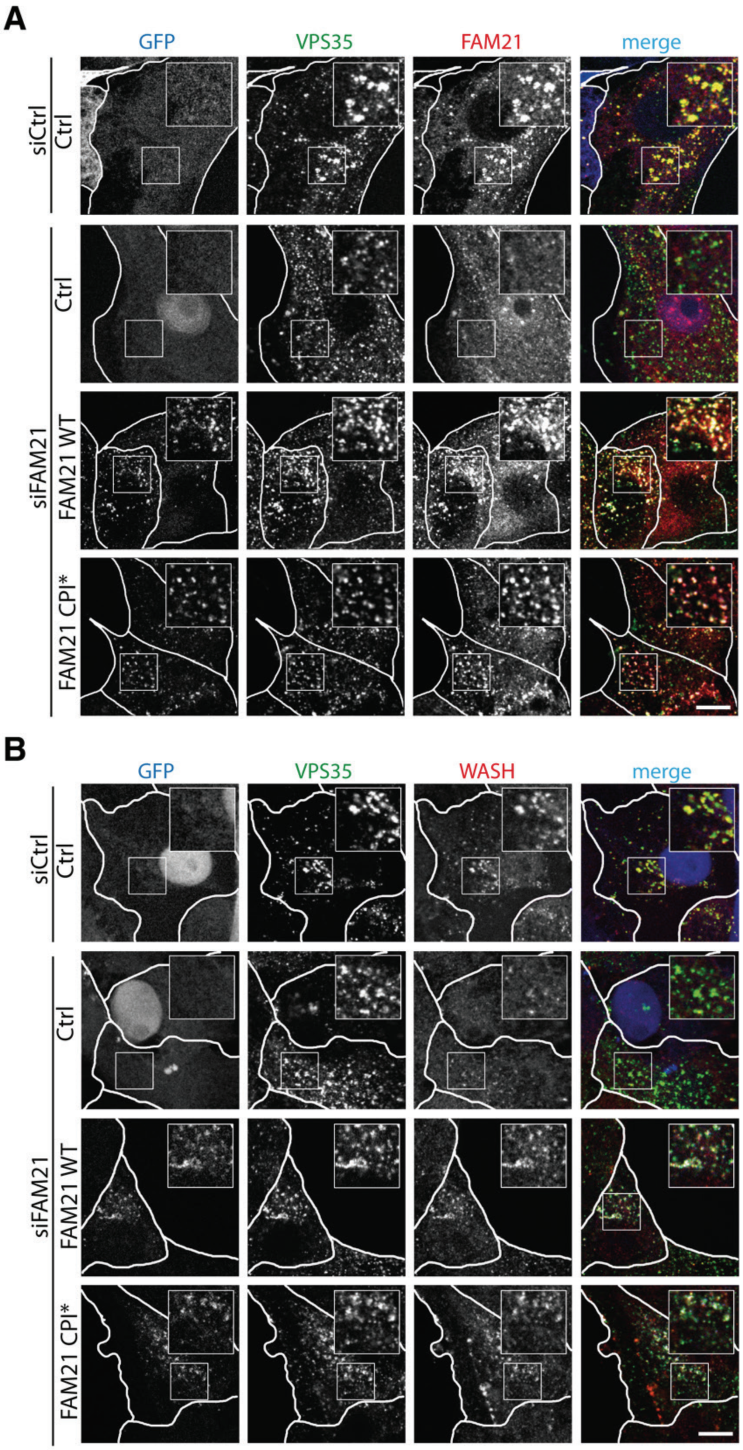


**Fig. S8. FAM21 CPI* reconstitutes a WASH complex that properly localizes at the surface of endosomes. A** FAM21 staining at the surface of endosomes is lost in FAM21 depleted cells and restored by GFP-FAM21 WT or CPI*. **B** WASH staining at the surface of endosomes is lost in FAM21 depleted cells and restored by GFP-FAM21 WT or CPI*. Scale bars: 10 µm.

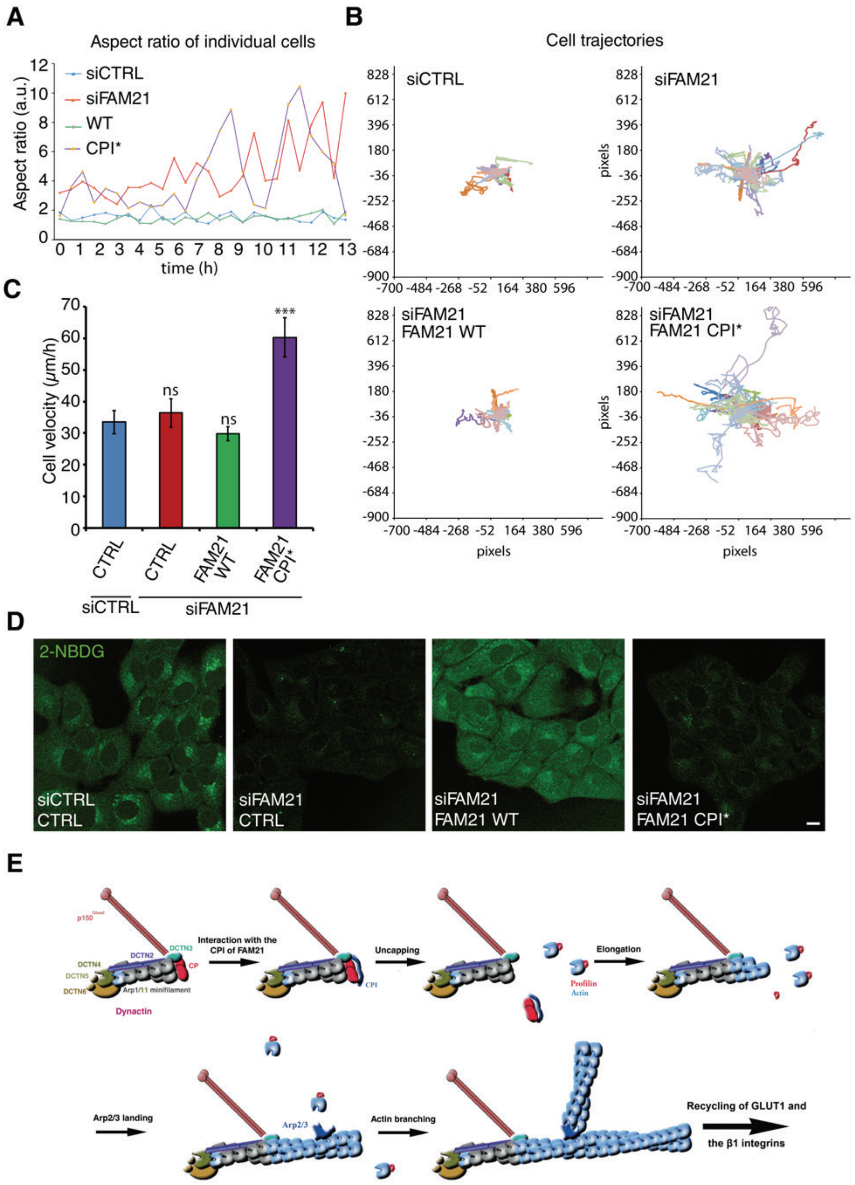


**Fig. S9. Characterization of FAM21 depleted cells restored with FAM21 CPI*. A-C** Cell migration of FAM21 depleted cells expressing FAM21 wild type, FAM21 CPI* or not. 20 cells per condition, 2 independent experiments. **A** Aspect ratio of individual cells from the Movie S5. **B** Trajectories of single migrating cells. **C** Average velocities. Mean ± s.e.m., one-way ANOVA, *** $P<0.0001$, ns: not significant. **D** Microscopic examination of 2-NBDG uptake. Quantification of intracellular fluorescence is displayed in Fig. 5E. Scale bar: 10 µm. **E** Model showing the cooperation between Dynactin, WASH and the Arp2/3 complex. The Arp1/11 minifilament of Dynactin is first uncapped by the CPI motif of FAM21. An actin filament can then be elongated in the presence of Profilin-actin. Finally, the VCA of WASH activates the Arp2/3 complex, which can land on the actin filament elongated from Dynactin. In the process, Dynactin becomes an integral component of the branched actin structure.

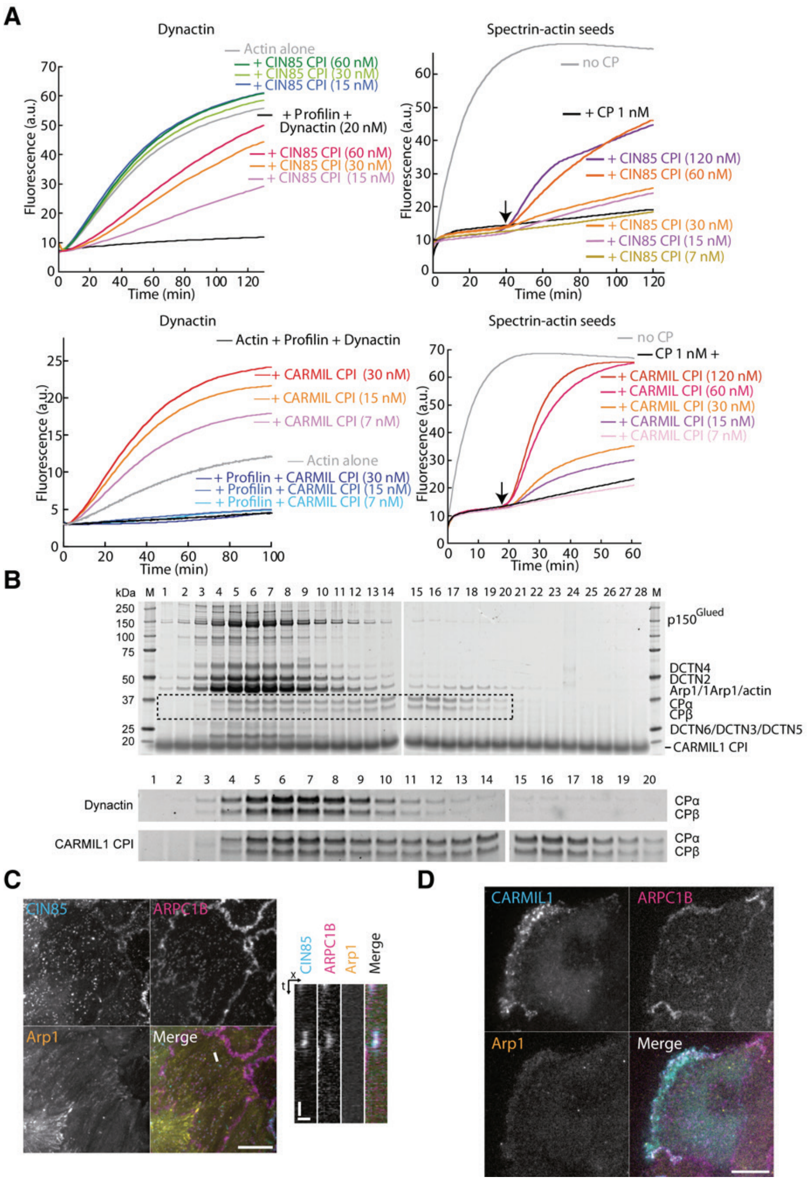


**Fig. S10. CPI motifs of CIN85 and CARMIL1 uncap Dynactin. A** The CPI motifs of CIN85 and of CARMIL1 are able to uncap Dynactin in vitro (left panels) and capped actin filaments (right panels). **B** Native Dynactin in gel filtration column in the presence of CARMIL1 CPI. The enlarged patterns show CP behavior in presence or absence of CARMIL1 CPI. **C** MCF10A cells expressing GFP-CIN85, mCherry-Arp1 and iRFP-ARPC1B were imaged live by TIRF microscopy. To enhance the detection of clathrin coated pits, the image corresponds to a time projection of 100 frames (1.5 s for a movie acquired at 0.67 Hz). The line corresponding to the kymograph is displayed on the merge image. The Arp2/3 complex and CIN85, but not Dynactin, colocalize at clathrin coated pits. Scale bar 10 µm and 1 µm /15 s for kymograph. **D** MCF10A cells expressing GFP-CARMIL1, mCherry-Arp1 and iRFP-ARPC1B were imaged live by TIRF microscopy. The Arp2/3 complex and CARMIL1, but not Dynactin, colocalize at the cell cortex. Scale bars: 10 µm.

### Captions for Movies

**Movie S1:** Time-lapse images of a filament growing under flow from an anchored Dynactin (green). Conditions: 1 µM actin (15% Alexa488 labeled, red), 2 nM Dynactin, 50 nM CPI, 1 µM Profilin. The image of GFP labeled Dynactin was acquired before the beginning of the time lapse, because of its fast bleaching. Alexa488 labeled filament was pseudo-colored in red. Scale bar: 2 µm. Corresponds to Fig.1G.

**Movie S2:** Fast live cell spinning disk confocal microscopy of a MCF10A cell line stably expressing GFP-ARPC5 (Arp2/3), mCherry-DCTN6 (Dynactin) and iRFP-WASH. Scale bar: 10 µm. Insets on the left, kymographs showing transient overlap, insets on the right, one endosome from the kymograph centered in the frame. The animation starts after 2 s. Corresponds to Fig.3B.

**Movie S3:** Single cell migration experiment of cells depleted of FAM21 and re-expressing iRFP-FAM21 WT, CPI* or not. 5 min frame rate. Scale bar: 10 µm.

**Movie S4:** Model of the interplay between Dynactin, the WASH and the Arp2/3 complex. The animation starts after 2 s. Corresponds to Fig. S9E.

**Movie S5:** MCF10A cells expressing GFP-CARMIL1, mCherry-Arp1 and iRFP-ARPC1B were imaged live by TIRF microscopy. CARMIL1 and ARPC1B colocalize at the cell cortex but do not colocalize with Arp1. Scale bar 10 µm. Corresponds to Fig. S10D.

**Movie S6:** MCF10A cells expressing GFP-CIN85, mCherry-Arp1 and iRFP-ARPC1B were imaged live by TIRF microscopy. Left panel shows the line corresponding to the kymograph on the merge image. CIN85 and ARPC1B colocalize, but do not colocalize with Arp1. Scale bar 10 µm and 1 µm /15 s for the kymograph. Corresponds to Fig. S10C.
